# Malaise trap samples of 1000 individuals per week suggest 4 million insects per hectare in the boreal zone

**DOI:** 10.64898/2026.06.05.730540

**Authors:** Luisa F. Rodriguez, Viktor Gårdman, Tomas Roslin, Otso Ovaskainen

## Abstract

A basic question in ecological research and biodiversity monitoring concerns the estimation of species abundances from trap catches. As a case in point, a Malaise trap can yield thousands of arthropod individuals, but how this count should be converted to numbers of individuals per unit area has remained an open question. Here, we supplement observational data with an experimental approach targeted at quantifying catchability. We released marked insects in a boreal forest and examined their capture rate by a grid of Malaise traps around the release location. We estimated insect movement rates, mortality rates, and Malaise trapping capture rates by fitting a joint species movement model to these data. As a methodological novelty, we show how to convert the movement model parameters to the expected number of captured individuals, given their actual population density. Our results show that multiplying the sample content by 30 000 yields a rough estimate of the number of individuals per hectare. This conversion factor depends on the species, generally decreasing with increasing body size. We apply the estimated conversion factors to conclude that typical boreal forest contains some four million insect individuals per hectare, out of which around half belong to Diptera.

**SIGNIFICANCE STATEMENT:** Traditionally, the Malaise trap method has been used for assessing the state of the local communities and to estimate population abundances. However, a topical question is: how does the number of individuals observed in a sample relate to the true density of individuals in the surrounding community? To answer this question, we implement a movement model parametrized by a carefully designed mark-recapture experiment, in which we are able to obtain taxon-specific conversion factors for different groups of insects. We found that different insect groups come with different conversion factors, causing a mismatch between sample contents and true community composition. Thus, treating the sample contents as a direct representation of the reference community will be misleading.

## INTRODUCTION

Monitoring biodiversity has never been more important, with studies revealing steep population declines around the world ^1,2^. With ongoing species loss, it is increasingly important that we monitor and track what we have left and what we are losing ^3,4^. Rational resource management and sound environmental policies rely on accurate estimates of population trends ^5,6^. Nonetheless, few if any studies have access to exhaustive counts of every individual. Instead, we are confined to samples obtained by different techniques.

Relating samples to estimates of species abundances is not straightforward, and remains one of the most basic challenges in ecological research and biodiversity monitoring ^7–10^. In organisms with discrete individuals, such as animals, species abundances are most naturally measured in units of population density, *i*.*e*., number of individuals per unit area. Estimating population densities has remained difficult for most organism groups, as abundance data are typically influenced not only by the actual population density, but also by the process by which the observations are made. This step is frequently omitted altogether, with samples taken to represent actual densities ^5,11–15^.

An urgent need for accurate estimates now relates to insect densities. Studies suggest a steep decline in insect diversity and biomass worldwide ^4,15,16^, prompting local ^17,18^, national ^19–21^ and global ^22^ efforts to estimate insect diversity and densities. Currently, the sampling method most frequently used to obtain a sample of the local community is the Malaise trap ^22–26^. Such a trap can sample as many as 750 arthropod individuals in a single day during ideal conditions ^27^. However, few studies have attempted to critically evaluate how the sample content relates to actual species densities (although see ^28,29^).

A Malaise trap is a passive intercept trap (Fig. 1). The insects entering the sample represent the set of insects flying into the vertical part of the trap, then continuing their path to the highest point of the trap and dropping into the collecting vial. Thus, the trap catch will be a compound function of flight paths crossing the trap, active avoidance by some agile insect groups, and the odds of entering the collecting vial after encountering the trap. Given these steps between actual population density and sample content, the abundances within a sample cannot be directly equated with insect densities in the surrounding landscape.

**Figure 1.**
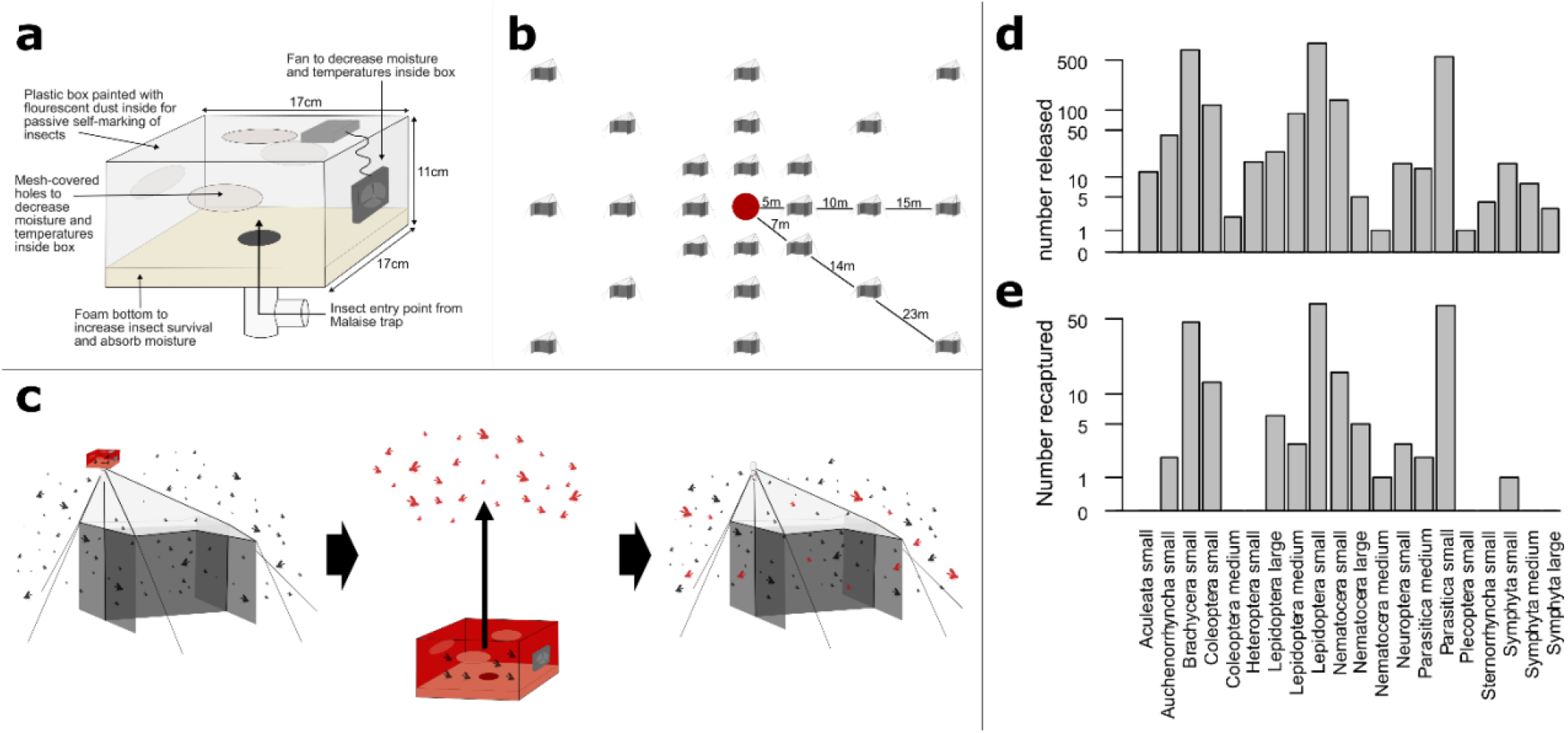
The mark-recapture experiment. The left-hand panel show (a) the design of the marking box for passive self-marking of insects, (b) the experimental design for the mark-recapture experiment, and (c) the workflow of sampling insects in dusted boxes for self-marking, releasing them once marked, and re-capturing the marked insect. The right-hand panels show the numbers of (d) released and (e) recaptured individuals for each species group and size class.

Mark-release-recapture studies provide a convenient way to relating sample contents to the latent reality ^30–33^. Targeted specifically at insect movements in heterogeneous landscapes, diffusion-advection-reaction models enable the estimation of movement, mortality and capture parameters based on data of where and when the marked individuals were released and recaptured ^34,35^. This approach has been further developed to include species traits in joint species movement models (JSMM), allowing to determine how species traits affect movement patterns and whether phylogenetically similar species share similar movement patterns ^36^.

In this study, we use Malaise trap data to show how samples (here: trap catches) can be converted to true densities by supplementing observational data with a mark-recapture experiment targeted at quantifying catchability. Using the joint species movement model implemented in the R-package Jsmm ^37^, we convert insect movement rates, mortality rates, and Malaise trapping capture rates to the expected number of captured individuals, given their actual population density.

## RESULTS

To estimate capture rates of boreal insects, we conducted a mark-recapture experiment (Fig. 1). In the experiment, we captured wild, forest-dwelling insects with Malaise traps, photographed the individuals for counting and taxonomic classification, marked them with fluorescent dusts, and released them within a network of Malaise traps in a boreal forest. Insects were released three days in a row using unique marking colour per day, followed by a four-day break. We then repeated this process three times, with a total of 13 release events (last event used only one colour). Marked individuals were recaptured within a 360 x 360 m grid of 24 Malaise traps around the release area. Traps were emptied daily during the release days and every second day during the four-day breaks in releases. The experiment ran for 24 days, recapturing 117 out of 2652 released individuals (Fig. 1).

To evaluate the relationship between population density and expected number of individuals captured by a Malaise trap, we analyzed the mark-release-recapture data with a model-based approach. In doing so, we assumed that the insects follow the classical diffusion-reaction model of movement ^38^. In this model, the probability density *u*(**x**, t) for a particular individual’s location ***x*** at time *t* evolves as

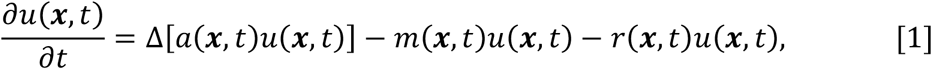

where the parameters relate to the rates of movement (*a*), mortality (*m*) and trapping (*r*). We note that including the trapping rate *r* in the reaction term corresponds to a cumulative capture process ^39^. This is appropriate for Malaise trapping, where captures are not instantaneous but accumulate over time. Given the vast number of boreal insect species, of which a large proportion is not even known to science ^25^, no experiment can be expected to capture the majority of species. For this reason, it is not feasible to estimate model parameters for each boreal insect species that may exist. Instead, as a pragmatic alternative, we grouped the released individuals in 20 groups based on their taxonomy and size (SI Appendix, Table S1), and assumed that the movement, mortality, and trapping parameters are group specific. We parameterized the diffusion-reaction model in the joint species movement modelling (JSMM) framework ^36^, which enables fitting the model simultaneously for multiple species (or species groups in our case) instead of fitting it separately for each species. Joint parameterization brings statistically efficiency, as it enables borrowing information among the species, leading to more accurate parameter estimation especially for those groups with limited numbers of recaptures ^36^. We fitted the JSMM model by applying the Bayesian parameter estimation scheme implemented in the R-package Jsmm ^37^.

Even if the number of recaptures was limited in our experiment, the data allowed for substantial learning of the parameters, as evidenced by the posterior distributions being different from and typically more concentrated than the prior distributions, and the parameter estimates differing among the groups of species (Fig. 2). The joint modelling framework makes it possible to ask how model parameters depend on properties of the species group, which we characterized by the size class and the taxonomic class (see Supplementary material Table 1). We found some level of statistical support for small insects moving at higher rate than large insects (posterior probability Pr=0.84), mortality rate being lower for large insects than for small insects (Pr=0.81), and recapture rate being higher for large insects than for small insects (Pr = 0.89).

**Figure 2.**
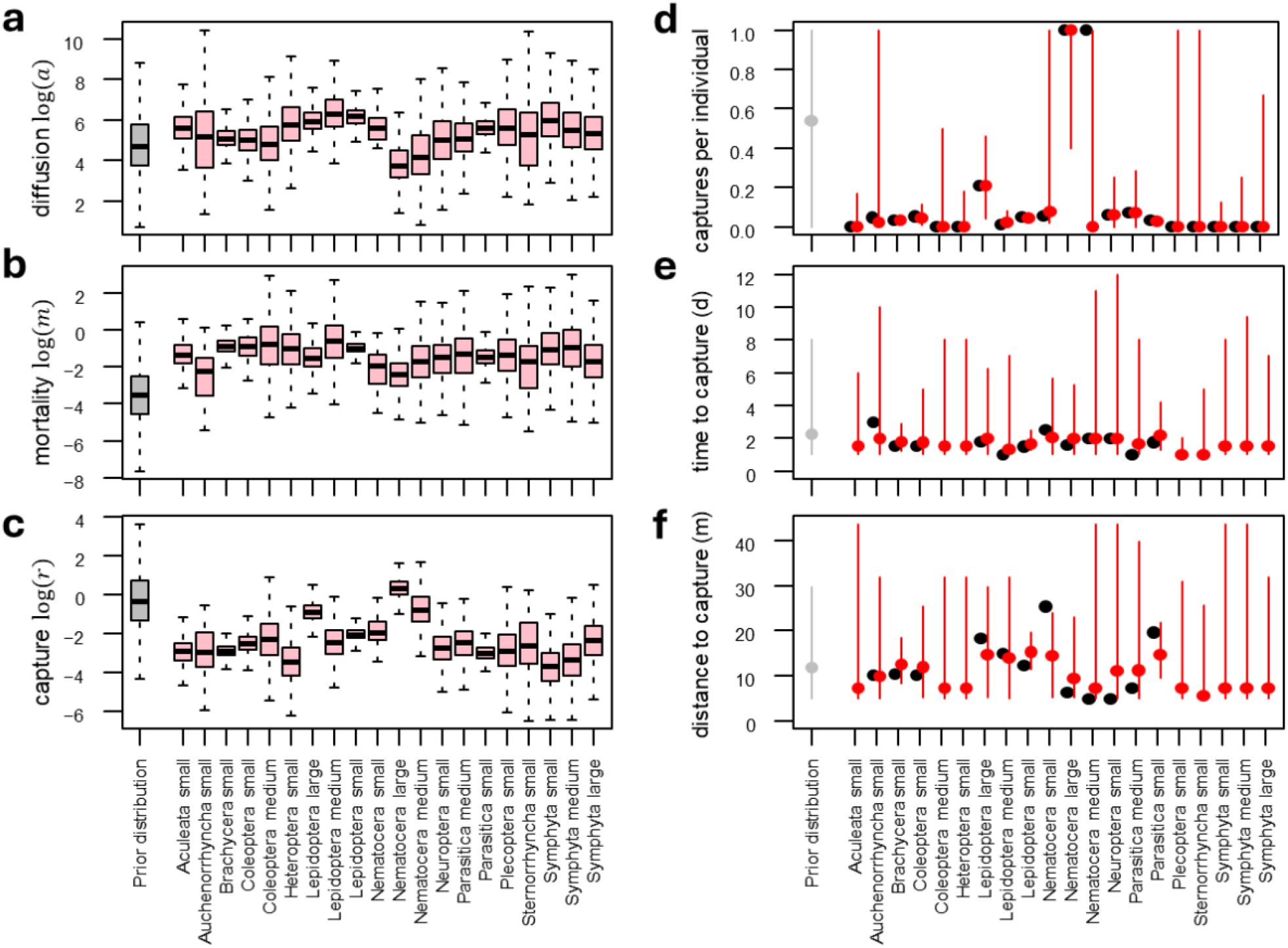
Joint species movement model parameterized with the mark-recapture data. The left-hand panels show posterior distributions of species-group specific parameters of (a) movement rate measured by diffusion parameter, (b) mortality rate, and capture rate (c). The right-hand panels evaluate model fit by comparing the observed data (black dots) to the predictive posterior distributions (red dots with error bars) in terms of (d) mean number of recaptures per released individual, (e) mean time between release and recapture, and (f) mean distance between release and recapture. Note that for several species groups the observed value is missing in panels (ef) due to lack of any recaptures. In all panels, the grey colours correspond to the prior distribution.

We evaluated model fit by comparing posterior and prior predictive data to the actual data in terms of the proportions of trapped individuals, the times between captures, and the distances between the captures (Fig. 2). Reflecting the limited numbers of recapture in our data, the posterior predictive distributions of these ecologically relevant summaries were associated with substantial uncertainty (Fig. 2). Yet they generally fitted the data much better than the prior predictive distributions (Fig. 2), suggesting the validity of the model structure and the parameterization.

As a key methodological novelty, we used the parameterized diffusion-reaction model to convert the actual abundance *Z* (number of individuals per hectare) into the expected number of individuals *N* that will accumulate in a Malaise trap during a given duration *T*. To implement such a conversion, we considered a hypothetical landscape consisting of a large rectangular domain of area *A*, with a single Malaise trap placed in the middle (Supplementary material Fig. 1). We further considered as the initial conditional a single individual that was located anywhere in the landscape, as modelled through the uniform distribution *u*(***x***, *t*_0_) = 1/*A*. By evaluating the diffusion model (Eq. 1) numerically over a trapping duration *T*, it is possible to compute the probability *p* by which the individual would become trapped according to the continuous capture process of Malaise trapping ^37,39^. Given an actual abundance *Z*, the expected number of individuals in the landscape will be *ZA*, and hence the expected number of trapped individuals will be E[*N*] = *ZAp*. Thus, *Z*^∗^ = 1/(*Ap*) is the population density that results in the expected number of trapped individuals being E[*N*] =1. This means that *Z*^∗^ is the conversion factor that we aimed to estimate, as multiplying the number of captured individuals by *Z*^∗^ yields an estimate of actual population density.

The posterior median estimates of actual abundances per trapped individual per day varied between 2000 (large Nematocera) and 100 000 (small Symphyta) (Fig. 3a). To predict population densities of boreal insects, we reanalyzed 299 samples that originated from 40 sites participating in the Global Malaise Trapping (GMT) project ^40,41^. Averaged over the samples, the mean number of captured individuals was 140 per day of sampling, whereas the mean estimated total insect density was 4.3 million individuals per hectare, and hence 430 individuals per square meter. There was expectedly major variation among the insect groups, with Plecoptera being least abundant (5600 individuals per hectare) and Nematocera (1.3 million individuals per hectare) and Brachycera (1.6 million individuals per hectare) the most abundant (Fig. 3b). Due to variation in group-specific conversion factors, the ratio between the estimated total insect abundance and total number of trapped individuals varied among the samples, yet the mean estimate of ca. 33 000 individuals per hectare per trapped individual per day remained relatively stable (Fig. 3c). Thanks to estimating the full posterior distribution of this estimate, we can quantify its uncertainty: while the posterior median estimate is 33 000 individuals per hectare per trapped individual per day, the interquartile range of this estimate is [27 000 … 40 000], and its 95% central posterior interval is [17 000 … 59 000].

**Figure 3.**
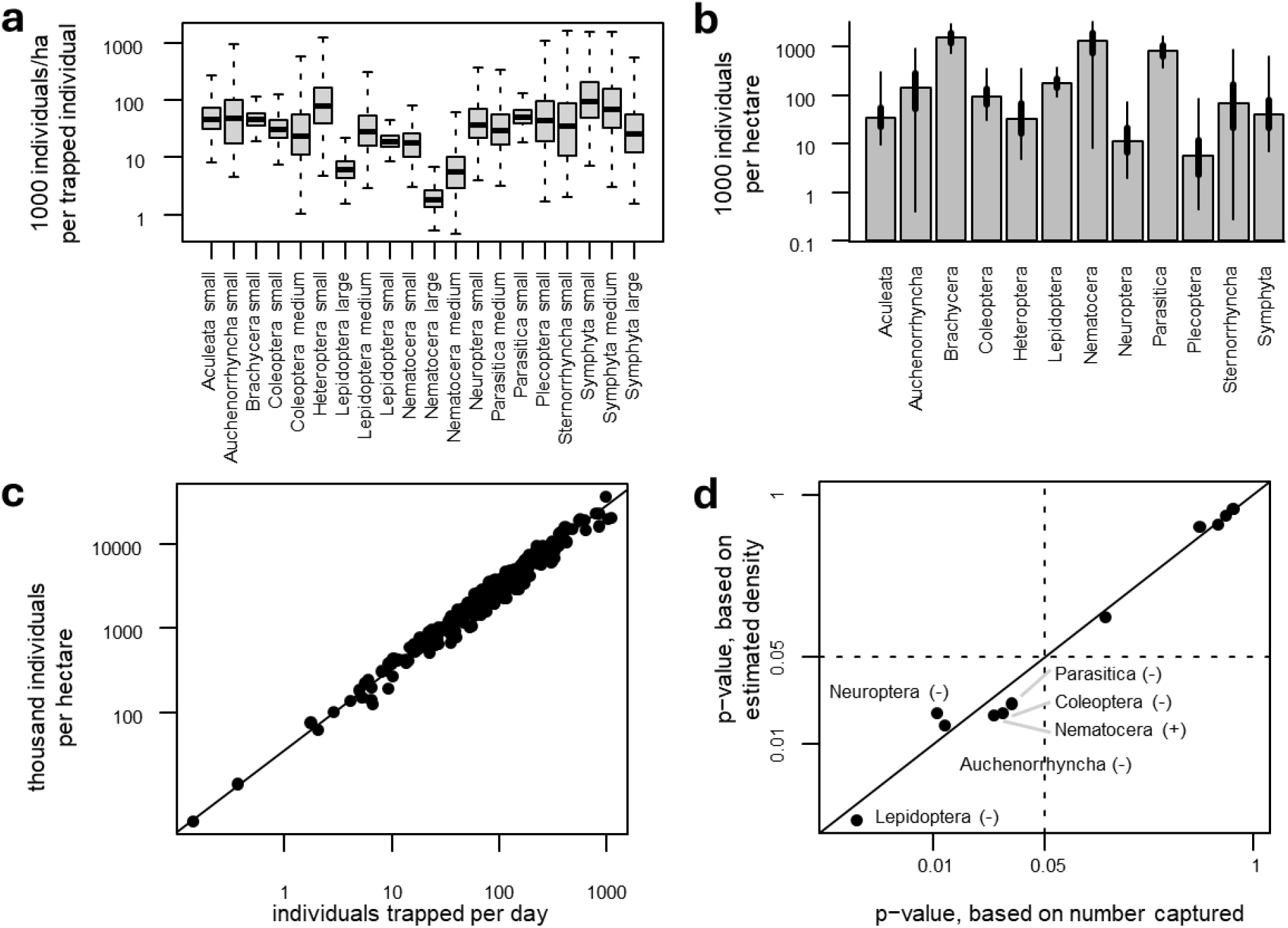
Estimated conversion factors and their use in ecological inference. Panel (a) shows the posterior distributions of the conversion factors *Z*^∗^ for different groups of insects, showing the expected actual population density (in units of thousand individuals per hectare) if one individual was captured during one day of Malaise trapping. Panel (b) shows the estimated population densities of different taxonomical groups, averaged over 299 Malaise trap samples obtained on 40 sites. The bar shows posterior median estimate, the thick line the posterior interquartile interval, and the thin line the posterior central 95% interval. Panel (c) shows, for each of the 299 Malaise traps samples, the estimated population density of all insects, as compared to the number of individuals captured per day of Malaise trapping. Panel (d) shows p-values from linear regression models where response is proportion of individuals belonging to each taxonomic group, with latitude as the sole explanatory variable. The p-value is computed either from a model where the proportion of individuals is counted from the numbers captured (x-axis) or from estimated population densities (y-axis). For statistically significant cases (p<0.05), we show the name of the taxonomical group and whether its proportion increases (+) or decreases (-) with latitude.

To illustrate how the estimated population densities may be propagated to ecological analyses, we asked how proportions of individuals belonging to taxonomical groups varied with latitude. We found the proportion to decrease with latitude in a statistically significant manner for Lepidoptera, Neuroptera, Auchenorrhyncha, Parasitica and Coleoptera, whereas the proportion increased only for Nematocera (Fig. 3d). While we recorded consistent results whether we computed the proportions based on the raw counts or the estimated densities, the latter yielded on average more significant relationships (Fig. 3d), suggesting that removing the effect of the observation process may lead to cleaner ecological relationships.

There were clear discrepancies in the relationship between sample composition and landscape composition (Fig 4). We found mosquitoes (Nematocera) to be overrepresented in malaise samples in relation to landscape abundances, composing 50 % of all insects in a malaise sample but only 31 % of landscape abundances. By comparison, the number of flies (Brachycera) and parasitoid wasps (Parasitica) were underestimated, with flies constituting 23 % of a malaise sample but 36 % of landscape abundances, and parasitoid wasps constituting 12 % of a malaise sample but 19 % of landscape abundances. These three taxa together make up roughly 86 % of a malaise sample and 86 % of landscape abundances, highlighting that these three taxa dominate the flying insect communities both in the traps and boreal forests.

**Figure 4.**
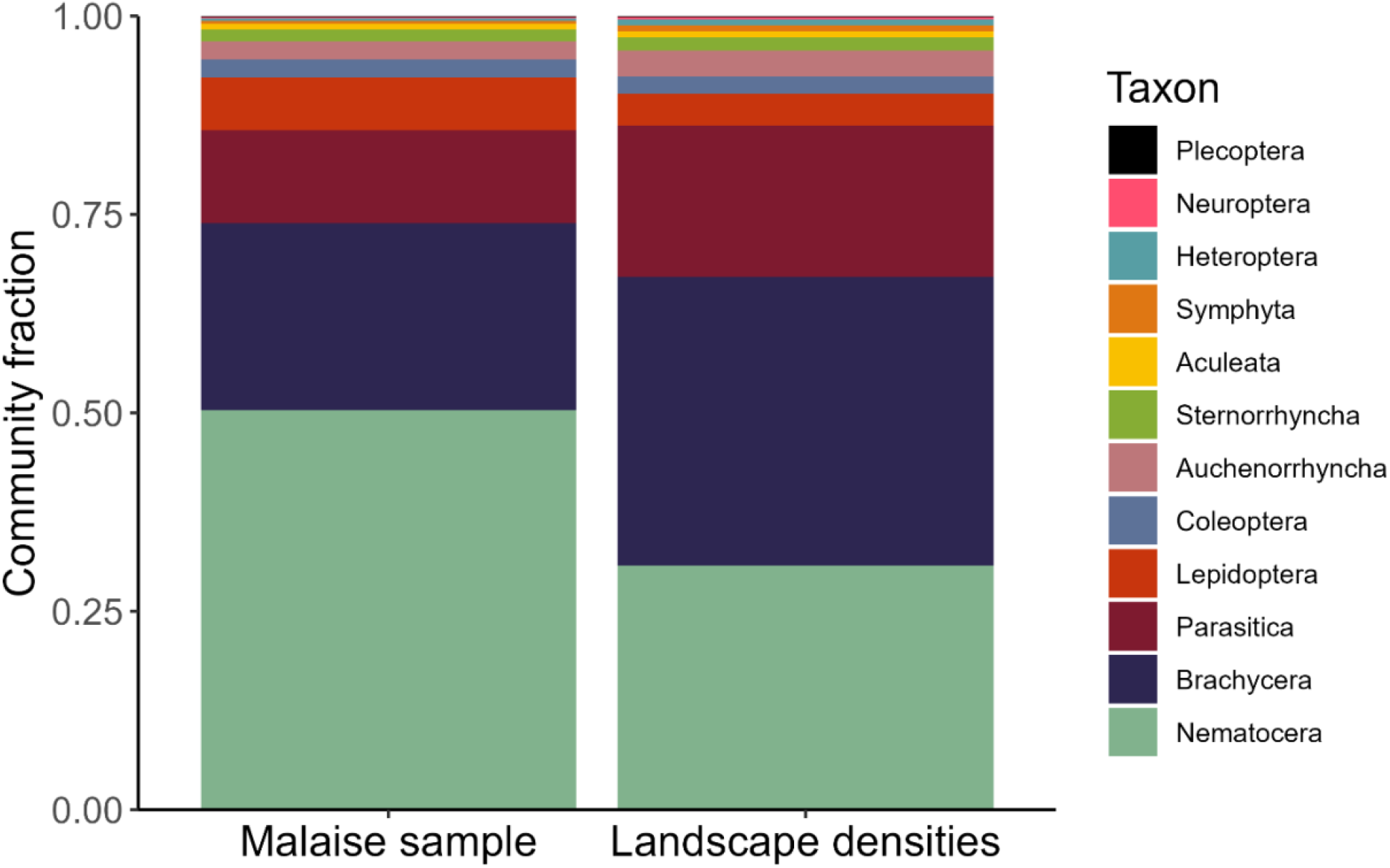
The relative abundance of taxa observed in a standard malaise sample from a boreal forest (left), and the corresponding true densities of taxa in the landscape (right). The proportion of landscape densities are based on posterior median estimates of the conversion rates shown in Fig. 3a.

### Conclusions

This study provides a new framework for converting counts observed in a sample to densities in the actual reference community. As a case study, we present a novel passive marking device for mark-release experiments of small insects in Malaise traps. Using the recapture rates from our experiment, we were able to estimate the true densities of insects in the surrounding landscape. We find that the taxon composition in a standard Malaise sample does not simply correspond to the true composition of flying insect abundances in the surrounding landscape. However, some broader patterns remain true; both Malaise samples and boreal forests are dominated by flies, mosquitoes, and parasitoid wasps. Using the mean conversion rate for an entire Malaise sample and a standard sample size of 140 individuals per day of sampling, we estimate that a hectare of boreal forest sustains roughly 4 300 000 insects. Thus, this study outlines an approach for converting sample contents to true, taxon-specific population densities in the reference community. While here applied to a specific type of sampling (Malaise traps), we stress that the general framework is applicable to other types of sampling.

## DISCUSSION

Most previous research on variation in animal abundances has been based on indices or proxies of population densities rather than on actual population density, compromising the accuracy and interpretability of the results achieved ^7–10,42,43^. To relate sample contents with surrounding densities, we need targeted experiments aimed at quantifying the probability of an individual reaching the sampling device and adding to the sample. Our case study shows that acquiring quantitative abundance estimation from large-scale observational data, supplemented by small-scale experimental data, is possible even for insects, as one of the most diverse and poorly known species groups in the world ^44,45^.

As a key outcome, we estimate that a mean density of 430 insect individuals passes by every square meter across a summer day in the boreal zone, or 4 300 000 individuals per hectare. The number is staggering, but we believe it to be well within reason. Malaise trapping and barcoding have revealed that insect densities and diversity vastly exceed those perceived by a passive human observed ^19,20,46^. Furthermore, the global consumption of insects by insectivorous birds in forests is estimated at 300 million tons per year ^47^. Adding predation by bats ^48^, spiders ^49^, and predacious insects ^50^, along with mortality caused by weather, parasitoids, and pathogens ^51^ (see also mortality rates estimated in Fig. 2), it becomes evident that millions of insects per day and hectare are needed to sustain boreal systems.

The conversion factors here estimated will also shed light on a concern frequently raised in studies based on Malaise traps. Given the hundreds of insect individuals caught in a Malaise trap per day, is there not a risk that we endanger the populations that we aim to monitor and protect? Here, our estimate of 30 000 insect individuals per hectare escaping the trap for every individual caught should still our concerns. A mortality rate of 0.03 permille will hardly make a dent in the population and will be negligible to compared to the natural mortality rates by predation and other natural causes.

In terms of the relative dominance of different taxa, our estimates suggest that direct records of sample contents may provide a biased view ^44^. Since different taxa come with different conversion factors (Fig. 3a), observing one large Nematocera (typically crane flies) in the catch will signal “only” a few thousand individuals in the surrounding hectare, whereas one small Symphyta (saw fly) in the sample corresponds to hundreds of thousands of individuals in the surrounding landscape. However, the dominance of mosquitoes (Nematocera), flies (Brachycera), and parasitoid wasps (Parasitica) seen in Malaise samples reflects their true dominance in the landscape. Together, they make up 86 % of the flying insect community in the boreal zone, with 2.9 million individual dipterans passing by each hectare of forest every day. Importantly though, the conversion factor differs among these groups. Mosquitoes will be less abundant in the landscape than a sample suggests, while flies and parasitoid wasps will be more abundant. Overall, Dipterans tend to exhibit the highest species richness of all insect orders in boreal systems ^46,52,53^, and Diptera is likely the most diversity insect order in the world ^46,54^. Given their abundance and diversity, dipterans are likely also important in sustaining boreal ecosystem function (as seen in the Arctic ^55^) and should receive more conservation attention ^56^. With this in mind, applying the conversion factors to samples becomes increasingly important. If we want to understand the ecological significance of the world’s most abundant insect taxa, we must first know just how abundant they are. Here, simply extrapolating the composition in Malaise samples would provide a biased view, whereas applying the conversion factors provides a more trustworthy picture of their true abundances.

Converting raw counts to estimates of actual abundances in the landscape increased statistical power for inferring the latitude-dependence of species composition. As assumed by previous work, we found that most insect taxa decrease in abundance with latitude ^53,57– 59^. The only group to increase was mosquitoes (Nematocera) – and observation in line with the notion that Dipteran diversity and abundance generally increase with increasing latitude ^53,58^. Due to the statistical advantages that comes with the use of the conversion factors, we recommend using actual abundances rather than raw trap catches when inferring ecological relationships from Malaise trap samples.

While this study provides an idea of the abundances of insects in boreal forests, we are most likely underestimating the true densities of *all* insects in the landscape. As an example, ground-dwelling, frugivorous, and saproxylic beetles are common in boreal forests ^57,60–62^, yet seldom sampled by Townes’ style Malaise traps ^27,63^ and not captured in this study. Our specific density estimates are thus applicable to flying insects commonly caught in Townes’ style Malaise traps, not of the entire community of insects in boreal forests.

## MATERIALS AND METHODS

### Mark-recapture experiment

#### Marking device

To quantify mark-release-recapture experiments, we designed a novel trapping device that passively marked insects in Malaise traps, allowing for bulk marking of insects. This design was built on previous approaches by Stern and Mueller ^64^, Culbert *et al*. ^65^; and Dickens and Brant ^66^. A 500 ml volume Wide-Mouth LDPE Bottles from Thermo Scientific (Rochester, USA) was cut in half and the top part, containing the trap attachment, was glued into a hole of the lid of a plastic box of 11x17x17 cm (Fig 1a). The walls and roof of the box were painted with a thin layer of fluorescent dust using a small brush. Three fluorescent dusts were used, using one colour per box: yellow dust (Product ID. 35898-2) from Partykungen (Gävle, Sweden) as well as one red dust (Product ID. TP-45) and one blue dust (Product ID. TP-49) from Radiant Color (Houthalen, Belgium). This made it possible for the insects to mark themselves passively by walking on, or flying by, the sides or roof of the traps. This approach followed the logic that insects would attach the marking medium to non-lethal part on their bodies such as legs, abdomens, or scutella, rather than having eyes, antennae or spiracles covered. To provide shade and hiding spots for the insects, we added pieces of egg cartons inside the boxes. To avoid insects being damaged when transported, we attached a foam-bottom to the boxes. Furthermore, to increase air circulation and thus keep the temperatures from reaching excessive levels inside the box, we cut holes on each side, as well as on the top, of the boxes of approximately 16 cm^2^, and covered these holes with a fine mesh. In another attempt to increase air circulation, we covered one of the mesh-holes with a 25 cm^2^ 5V computer fan (Product ID EE40100S2-1000U-999) from Sunonwealth Electric Machine Industry Co., Ltd (Kaohsiung, Taiwan). The fan was connected to a battery holder glued onto the outside of the box. The fan created a wind funnel, decreasing temperature within the box on warm days. The decrease in temperature should not only promote survival but also help calm the insects ^67^. The boxes were screwed onto the Malaise traps using the upper bottle attachments in the lids, and thus functioned as inverted Malaise trap bottles, with the insects ending up in the fluorescent dust-covered boxes instead of ethanol. Note that this approach is not possible for all models of Malaise traps, as they need to have an attachment point for bottles on top of the trap. The traps used in this study were ez-Malaise Trap II, Townes Style (Product ID. BT1012) from MegaView Science Co., Ltd (Taichung, Taiwan).

#### Mark-recapture experiment

To obtain a natural composition of species for the mark-recapture trials, we set up 16 Malaise traps with the marking boxes next to forest edges and large bushes around the premise of The Swedish University of Agricultural Sciences in Uppsala (Lat 59.817829, long 17.655387), and an additional eight inside a boreal forest near the experimental area (Lat 60.025041, long 17.749144). The marked boxes were attached between 8:00 and 9:30 in the morning and collected after 24 hours, attaching new boxes at the same time for further collection the following day. We deemed that 24 hours would allow for a substantial number of insects to be captured without infringing on their health.

To obtain recapture rate of the marked insects, we used a boreal forest site near Vattholma in Sweden, (Lat 60.0248, long 17.7513). The site was dominated by 40–60-year-old pines. The understory was also represented by pine, with scattered findings of spruce. The forest floor was dominated by wild strawberries (*Fragaria vesca*), horsetails (*Equisetaceae*), and graminoids. A tarmac road passed 20 meters south of the site. In the forest, we set up a release point for marked insects. Eight Malaise traps were then erected in a square around a release point, of which the horizontal and vertical traps were five meters from the release point and the diagonal traps thus were set seven meters from the release point. Another square of eight traps was then added so that each horizontal and vertical trap was ten meters from the first square traps (fourteen meters for diagonal traps), and a final square of eight traps were set fifteen meters from the second square traps (23 meters for diagonal traps). In total, 24 four traps were used encompassing 129 600 m^2^ (Fig 1b). Insects were collected from the marking box Malaise traps and brought to the release point in the middle of the trapping area. The insects were gently poured from the boxes into a plastic bag using a funnel. To avoid crushing the insects, the plastic bags had cardboard to the inner sides, keeping the bags three-dimensional, and a white bottom. By photographing the content of each plastic bag before releasing the insects, we could later count the releases and divide the insects into suitable taxa. After acquiring the images, the insects were poured onto open petri dishes. To ensure the insects would not be hurt or escape in distress from the area upon release, we took two precautions. Firstly, we placed buckets upside-down on top of each petri dish, with the bottoms replaced with metal meshes and covered with hay. This provided a shaded, protected environment for the insects. Secondly, a tarpaulin two meters above the buckets was erected to protect the emerging insects from rain and provide them with shade. To be able to exclude individuals that died on the petri dish from the analyses, the petri dishes were removed 24 hours after release. Any insects remaining on the petri dishes were recorded by images, identified, and later removed from the total counts of released insects per taxon.

Thirteen daily catches of marked insect individuals were released at the site between the 26^th^ of June to the 21^st^ of July in 2022. We used one fluorescent dust for marking per day for three consecutive days. To limit the risk of mixing findings of individuals of the same colour released on different occasions, we waited four days after a batch of insects marked which each colour dust had been released before we released new individuals again. We emptied the traps daily during the release period and switched to every second day during the subsequent four days.

To count marked individuals in the recaptures, the content of each Malaise bottle was poured onto a tray and scanned under UV-light with a microscope under complete darkness. Each marked individual was identified to a minimum taxonomic level of Order. To account for differences in life history and physical build within Orders, we split individuals of Hymenoptera into *Symphyta* (Sawflies), *Aculeata* (wasps and bees) and *Parasitica* (Parasitoid wasps). Using the same logic, dipterans were divided into *Brachycera* (Flies) and *Nematocera* (Mosquitoes). To infer how many days the insects had been active in the release area before recapture, we noted the colour of the marking dust. Each individual was then photographed in ethanol next to a square with an area of 25 mm^2^, using a Nikon D7000 with a 18 – 55 mm objective, ISO 2000, F14, and 1/100 shutter. The insects’ areas were calculated by covering them in masks using TORAS ^68^ and relating the size of the masks to the known 25 mm^2^ squares. This method followed an approach similar to that used to obtain insect area measurements in the Bioscan-5M dataset ^23^.

To make sure that the fluorescent colour would be visible after the insects were submerged in ethanol, we conducted trials with wild individuals caught in the marking-boxes. Wild insects dusted with any of the three colours, as well as a known number of unmarked individuals, were poured alive into either of four bottles with 95 % ethanol and shaken. The dusting remained visible under UV-light on all marked insects after 72 hours in ethanol, irrespective of dusting colour. Only once did an unmarked individual become secondarily marked inside the bottles.

#### Size distribution of released insects

To estimate the size distribution of the released insects, we obtained size distributions of insect taxa in boreal forests from the Bioscan-5M dataset ^23^. In short, we filtered this dataset to only include findings between latitudes of 55.3 and 69.1 (representing the boreal zone) and divided the remaining findings into the taxa used in the mark-recapture experiment. We split the total span of sizes per taxon into three equally sized groups and assigned all individuals that fell within the lower 33% of the total size span as small, the middle 33 % as medium, and the upper 33 % as large. The ratios of individuals within each size class were then applied on our released individuals to estimate the number of released individuals per taxon and size class. To obtain estimates of the number of individuals within each taxon and size class captured in a weekly Malaise sample in the boreal zone, we used the data from the Global Malaise Trap Project (GMP) ^40,41^. We included 299 samples that originated from 40 sites from GMT within the boreal zone and again divided taxa into the three size classes using the ratios from the Bioscan-5M dataset. To be able to include insects not released or recaptured in the experiment, we calculated size classes for all taxa available in the Bioscan-5M dataset.

### Statistical analyses

All statistical analysis were conducted in R Version 4.5.3 ^69^. We applied the JSMM framework of Ovaskainen *et al*. ^36^ implemented in the R-package Jsmm ^37^. As initial setup for the Jsmm model m, we configurated the spatial information regarding the boreal forest in which the experiment was conducted and the capture-release sites (Fig 1b) as the domain named list (SI Appendix Fig. S1). Following the experimental design, we set the observation model as a continuous capture process (CCP) within the observation_effort named list composed of 576 and 13 capture and release events, respectively, and the experiment length has been set as 24 days (t=24) (Supplementary material Fig. 2). The trait information has been modelled by the model formula ∼*taxon* + *size* (Supplementary material Table 1), and no phylogenetic relationships have been assumed between the 20 groups. As process model, we assumed spatially explicit diffusion supplemented by mortality. Within the Jsmm framework, the movement and observation parameters are combined and log-transformed to enable multivariate normal model for *θ*_*s*_ for each group *s* (SI Appendix Table S2) to the vector:

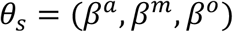

were *β*^*a*^ is the diffusion coefficient (unit *m*^2^/day, measuring the movement rate), *β*^*m*^ is the mortality rate (unit/day), and *β*^*o*^ is the capture rate defined within each of the capture sites. No dependence on spatial, temporal or spatio-temporal variables was assumed for any of the model parameters.

We assumed the default prior parameters, which correspond to mortality rate that leads to probability of death 0.5 during the study period; diffusion rate that leads to expected dispersal distance from birth to death being half the size of the study domain; and trapping rate that leads to capture probability 0.5 during the study period.

We sampled the posterior distribution of the model with the Markov chain Monte Carlo (MCMC) method of Rodriguez and Ovaskainen ^37^ with five chains, each of which we run for 1875 iterations, of which we considered the first 625 iterations as transient. We thinned the remaining samples by 5 to yield 250 posterior samples per chain and thus 1250 posterior samples in total. We examined MCMC convergence by computing the potential scale reduction factors of the model parameters (see SI Appendix Table S2).

## ACKNOWLEDGEMENTS

LFR was funded by the University of Helsinki-HIIT PhD Fellowship on multidisciplinary applications of AI. OO was funded by the Research Council of Finland (grant no. 336212 and 345110). OO and TR were funded by the European Research Council (ERC) under the European Union’s Horizon 2020 research and innovation programme (grant agreement No 856506: ERC-synergy project LIFEPLAN).

