## Supporting Information for "Malaise trap samples of 1000 individuals per week suggest 4 million insects per hectare in the boreal zone"

**This PDF includes:**

Figures SI to S3

Tables SI to S3

SI References

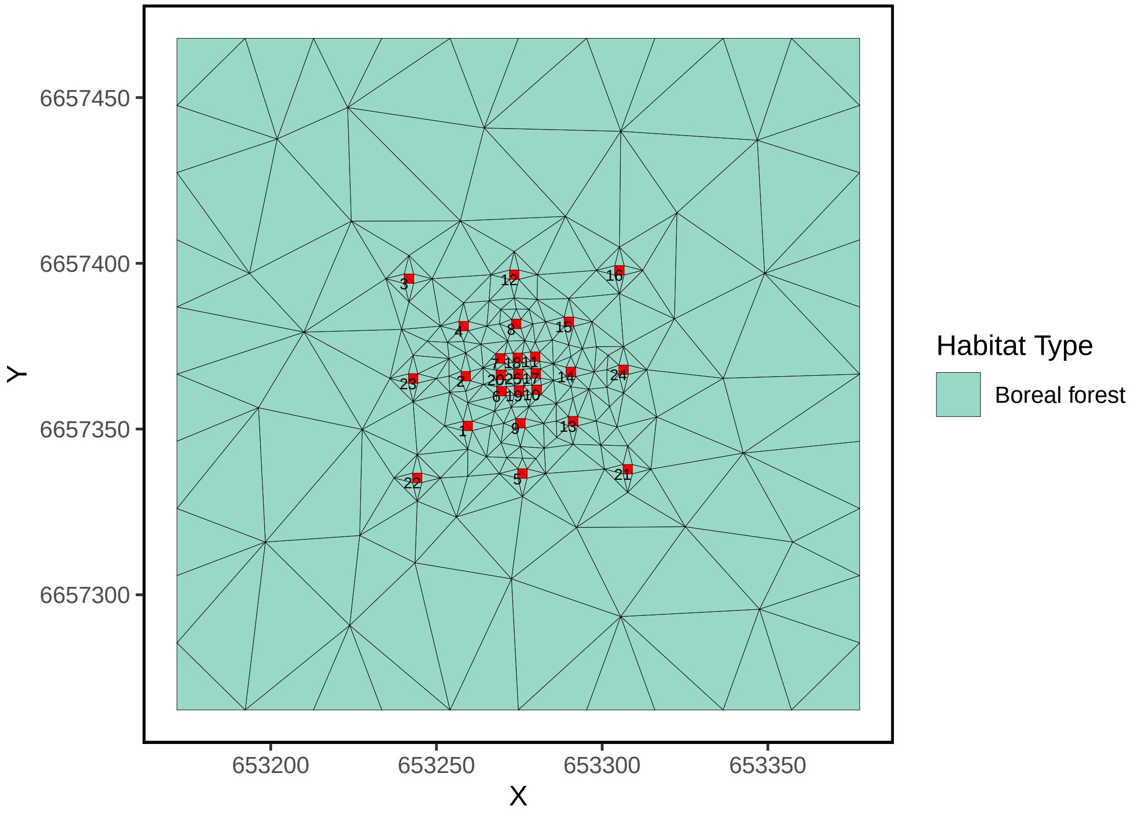


Figure S1: Map of the boreal forest study site located in Vattholma, Sweden (Lat 60.0248, long 17.7513) showing the position of the Malaise traps (red squares with labels 1 to 24, each with an area of 8 m2), the release site corresponds to the central square labeled as 25. The triangulation shown was used to implement the Joint species movement modelling framework [4] within the Jsmm R-package [3]. The rectangular domain has a total area of 41741.07 m2, and is composed of 490 elements and 266 nodes.

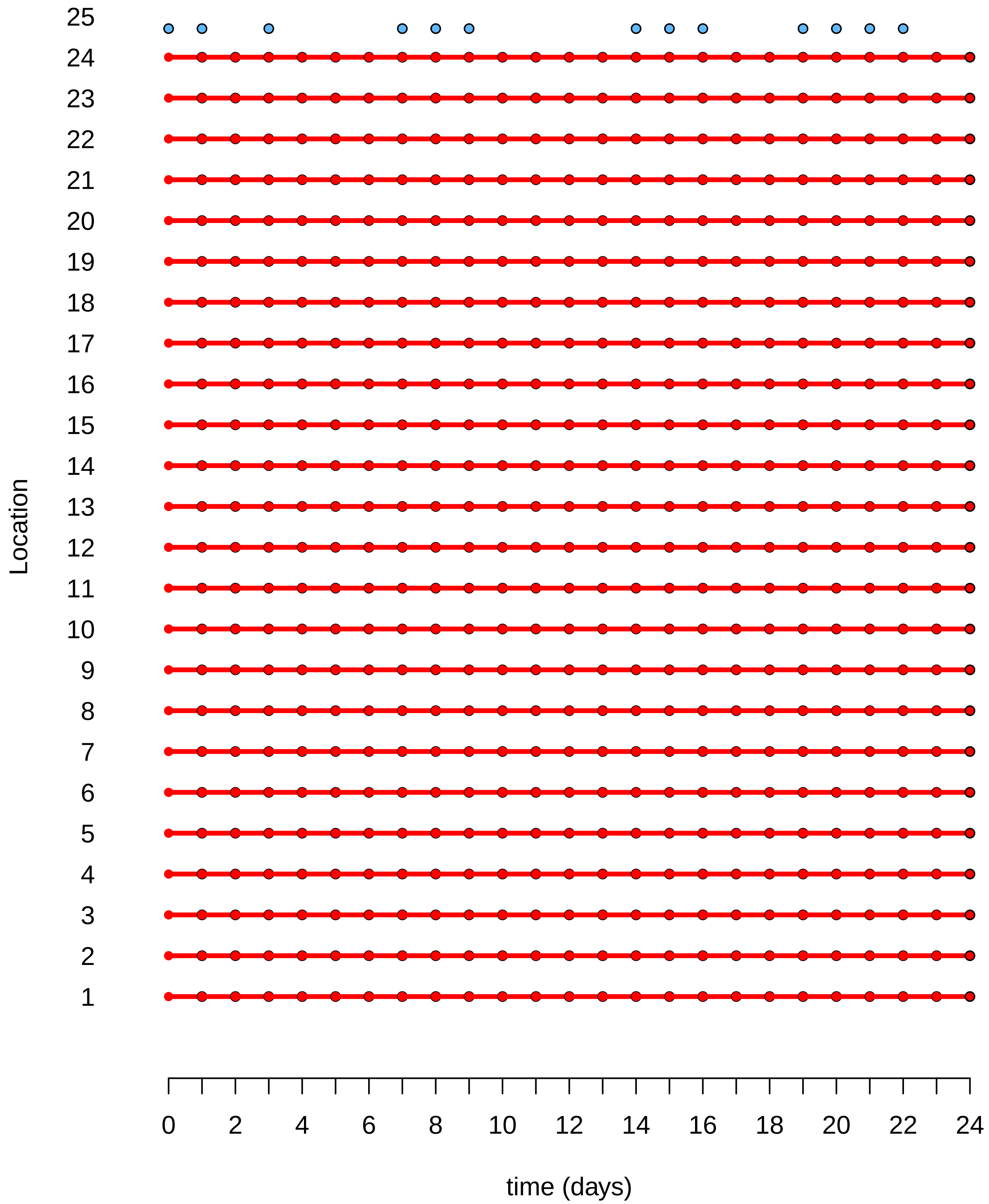


Figure S2: Observation effort scheme for CPP capture process within the Jsmm R-package [3]. Times in which releases and captures are possible are represented by blue, and red colors, respectively. The observation effort named list is composed of 576 and 13 capture and release events, respectively.

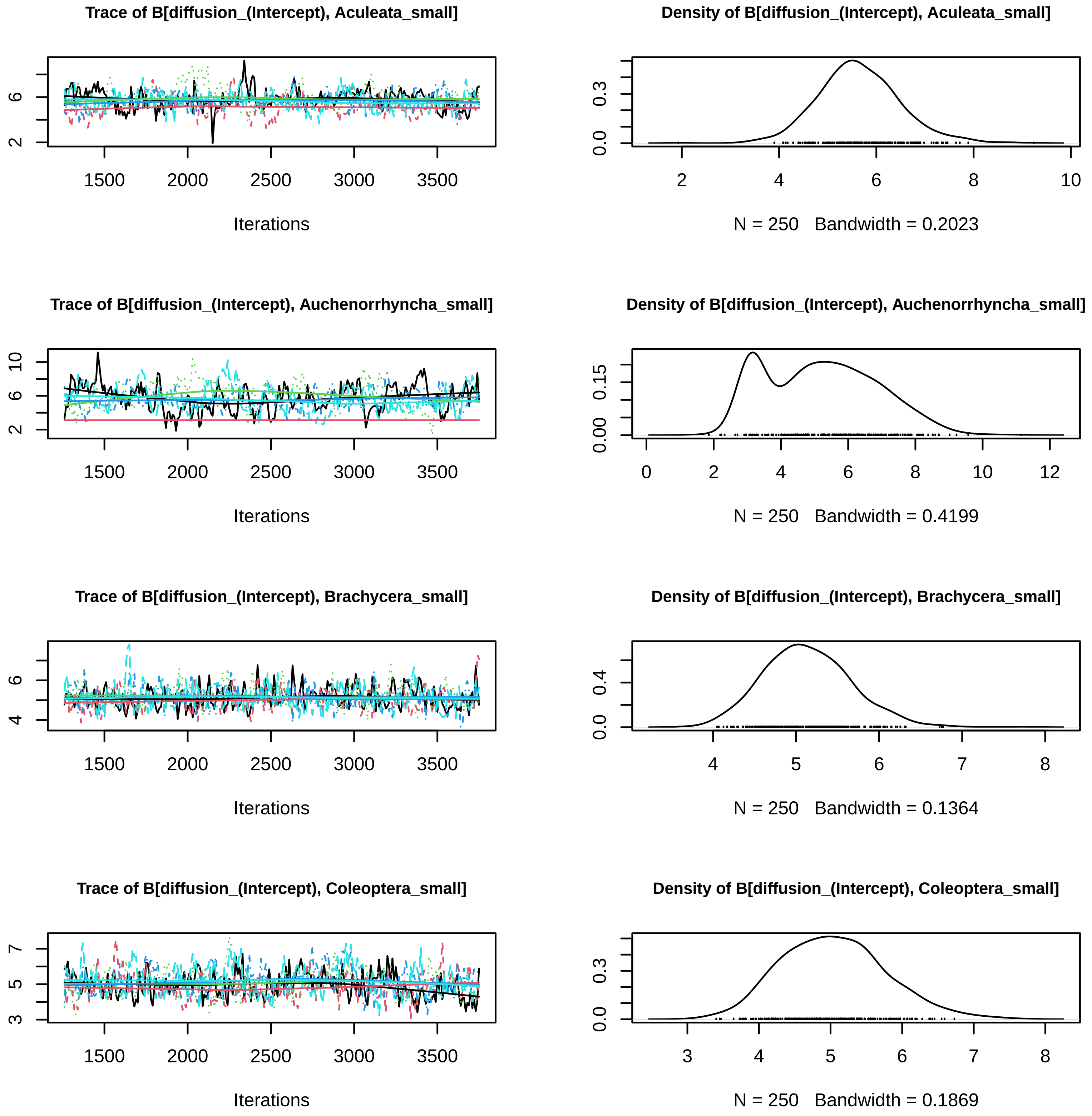
Trace of B[diffusion_(Intercept), Aculeata_small]
Iterations
Density of B[diffusion_(Intercept), Aculeata_small]
N = 250 Bandwidth = 0.2023
Trace of B[diffusion_(Intercept), Auchenorrhyncha_small]
Iterations
Density of B[diffusion_(Intercept), Auchenorrhyncha_small]
N = 250 Bandwidth = 0.4199
Trace of B[diffusion_(Intercept), Brachycera_small]
Iterations
Density of B[diffusion_(Intercept), Brachycera_small]
N = 250 Bandwidth = 0.1364
Trace of B[diffusion_(Intercept), Coleoptera_small]
Iterations
Density of B[diffusion_(Intercept), Coleoptera_small]
N = 250 Bandwidth = 0.1869

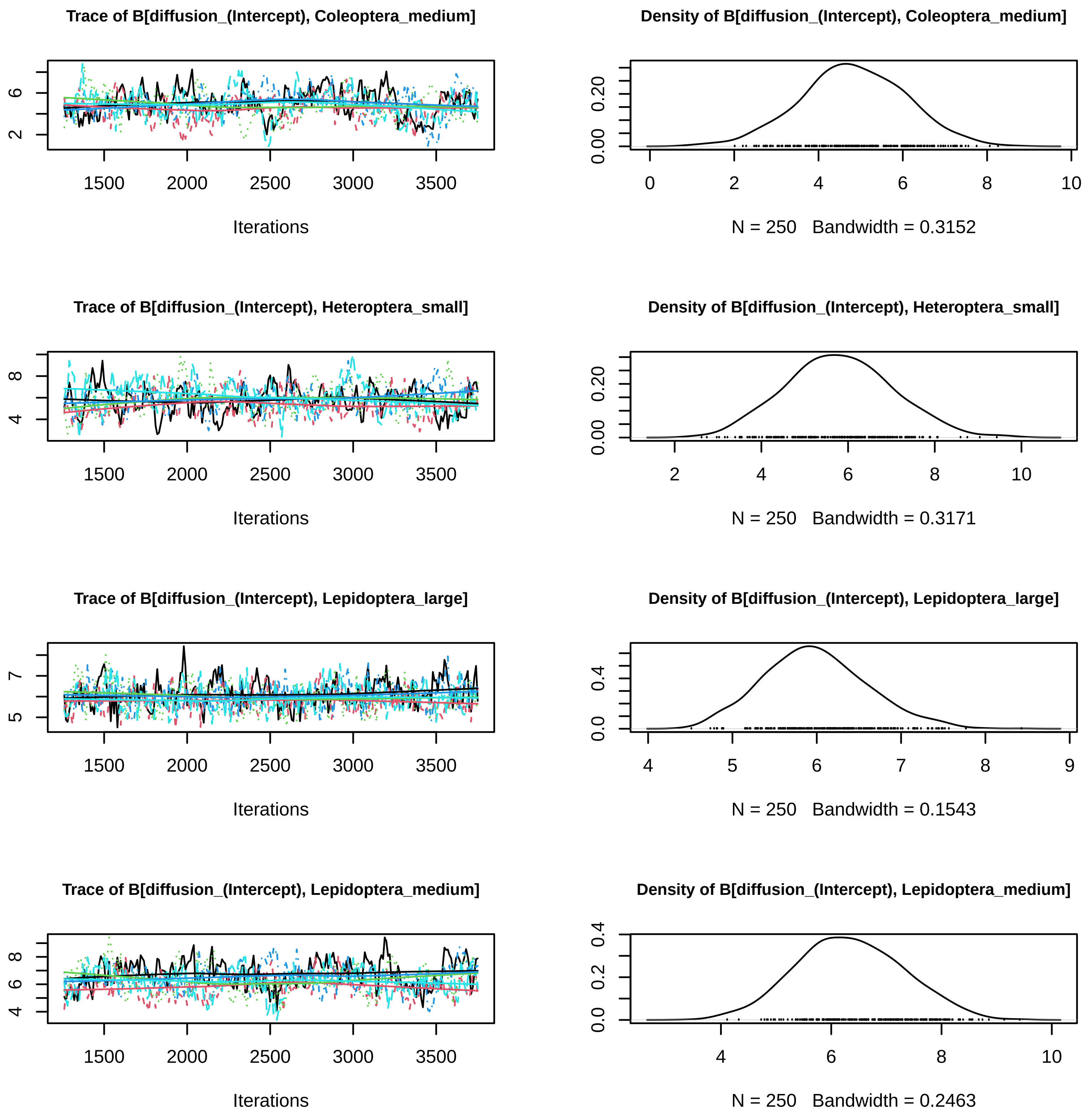
Trace of B[diffusion_(Intercept), Coleoptera_medium]
Density of B[diffusion_(Intercept), Coleoptera_medium]
N = 250 Bandwidth = 0.3152
Trace of B[diffusion_(Intercept), Heteroptera_small]
Density of B[diffusion_(Intercept), Heteroptera_small]
N = 250 Bandwidth = 0.3171
Trace of B[diffusion_(Intercept), Lepidoptera_large]
Density of B[diffusion_(Intercept), Lepidoptera_large]
N = 250 Bandwidth = 0.1543
Trace of B[diffusion_(Intercept), Lepidoptera_medium]
Density of B[diffusion_(Intercept), Lepidoptera_medium]
N = 250 Bandwidth = 0.2463
Iterations

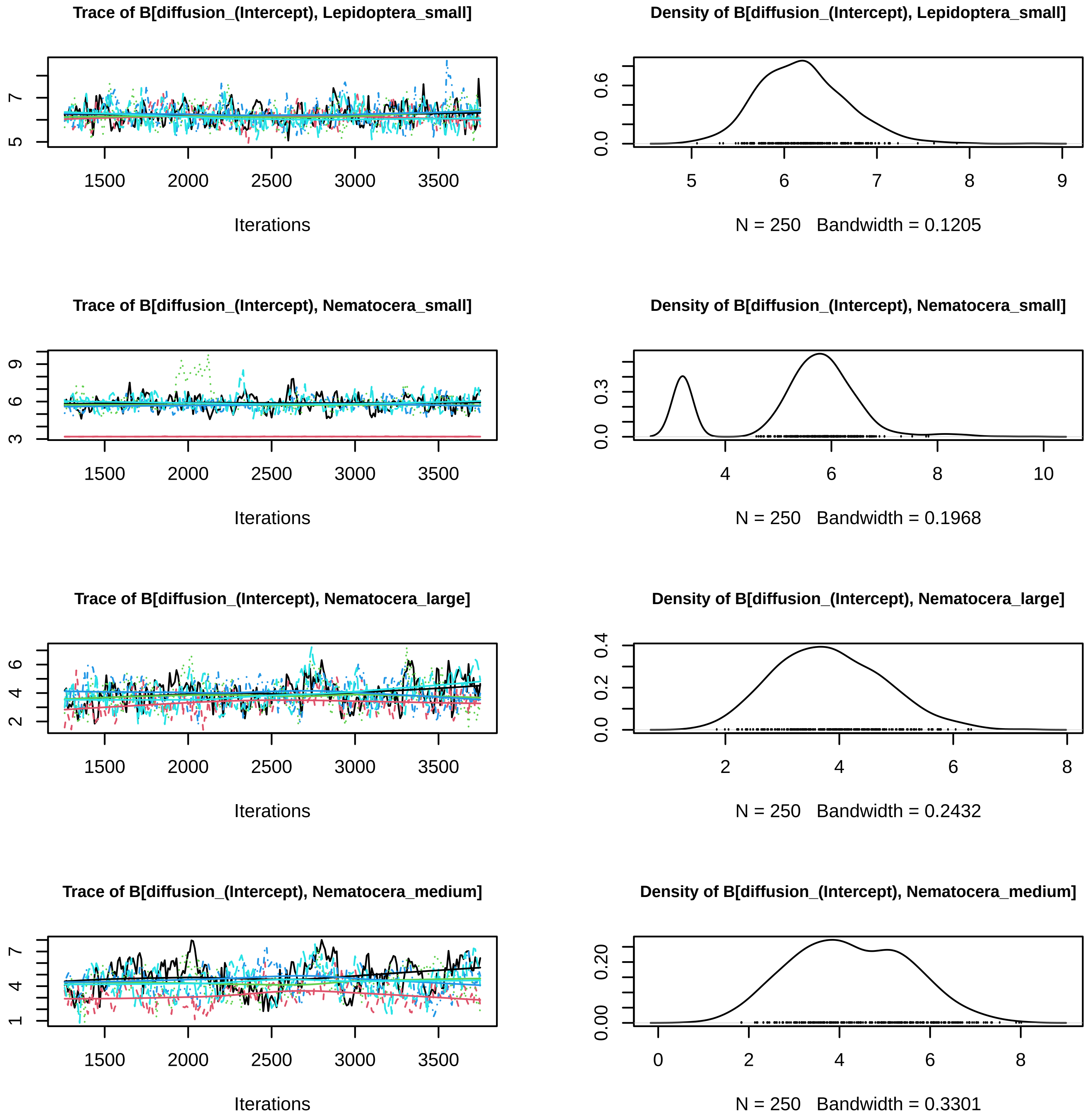
Trace of B[diffusion_(Intercept), Lepidoptera_small]
Iterations
Density of B[diffusion_(Intercept), Lepidoptera_small]
N = 250 Bandwidth = 0.1205
Trace of B[diffusion_(Intercept), Nematocera_small]
Iterations
Density of B[diffusion_(Intercept), Nematocera_small]
N = 250 Bandwidth = 0.1968
Trace of B[diffusion_(Intercept), Nematocera_large]
Iterations
Density of B[diffusion_(Intercept), Nematocera_large]
N = 250 Bandwidth = 0.2432
Trace of B[diffusion_(Intercept), Nematocera_medium]
Iterations
Density of B[diffusion_(Intercept), Nematocera_medium]
N = 250 Bandwidth = 0.3301

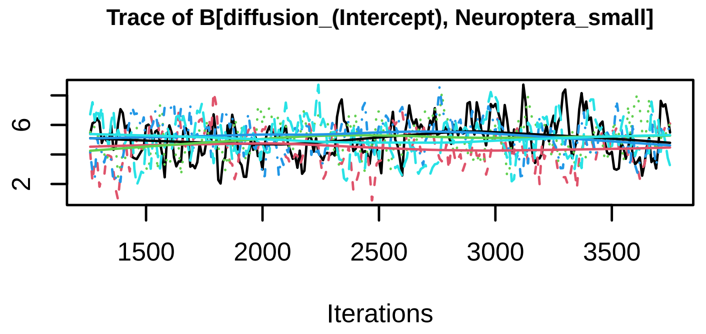
Trace of B[diffusion_(Intercept), Neuroptera_small]
2
6
1500
2000
2500
3000
3500
Iterations

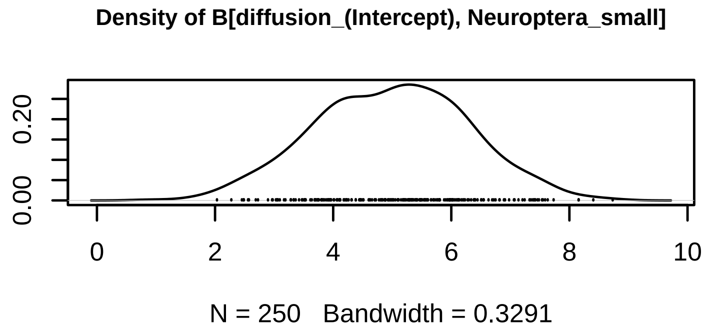
Density of B[diffusion_(Intercept), Neuroptera_small]
0.00
0.20
0
2
4
6
8
10
N = 250   Bandwidth = 0.3291

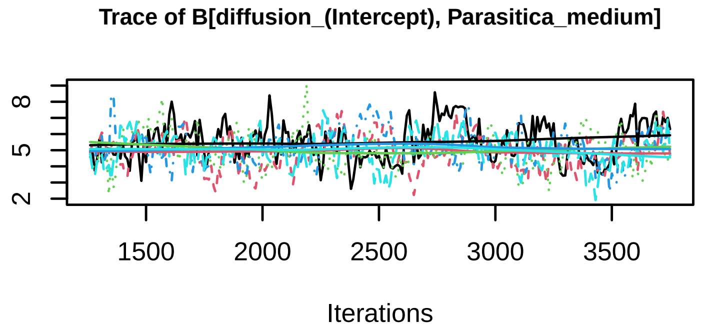
Trace of B[diffusion_(Intercept), Parasitica_medium]
2
5
8
1500
2000
2500
3000
3500
Iterations

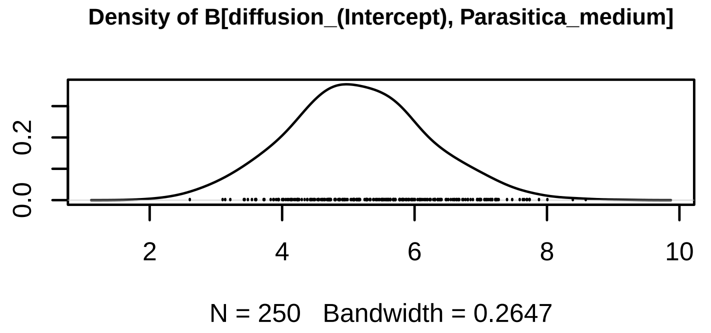
Density of B[diffusion_(Intercept), Parasitica_medium]
0.0
0.2
2
4
6
8
10
N = 250   Bandwidth = 0.2647

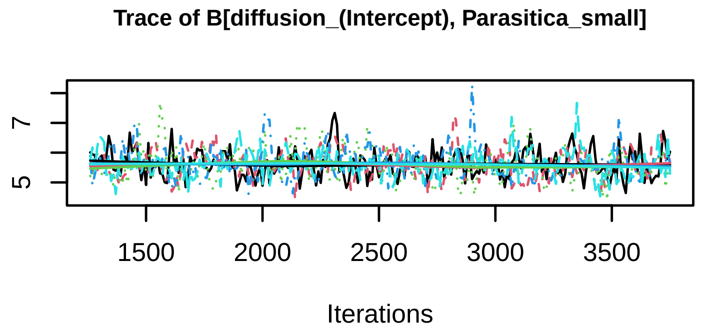
Trace of B[diffusion_(Intercept), Parasitica_small]
5
7
1500
2000
2500
3000
3500
Iterations

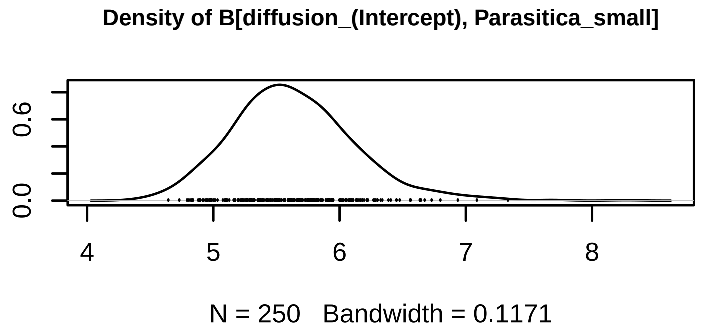
Density of B[diffusion_(Intercept), Parasitica_small]
0.0
0.6
4
5
6
7
8
N = 250   Bandwidth = 0.1171

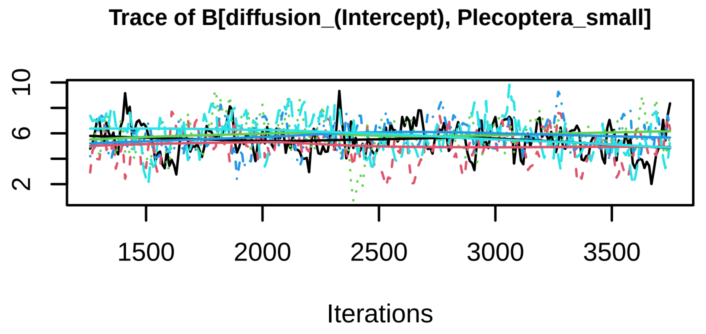
Trace of B[diffusion_(Intercept), Plecoptera_small]
2
6
10
1500
2000
2500
3000
3500
Iterations

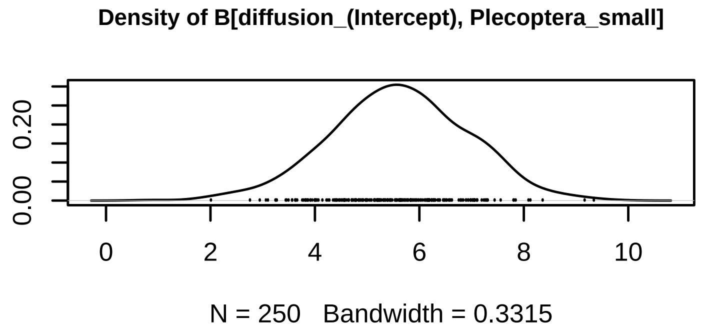
Density of B[diffusion_(Intercept), Plecoptera_small]
0.00
0.20
0
2
4
6
8
10
N = 250   Bandwidth = 0.3315

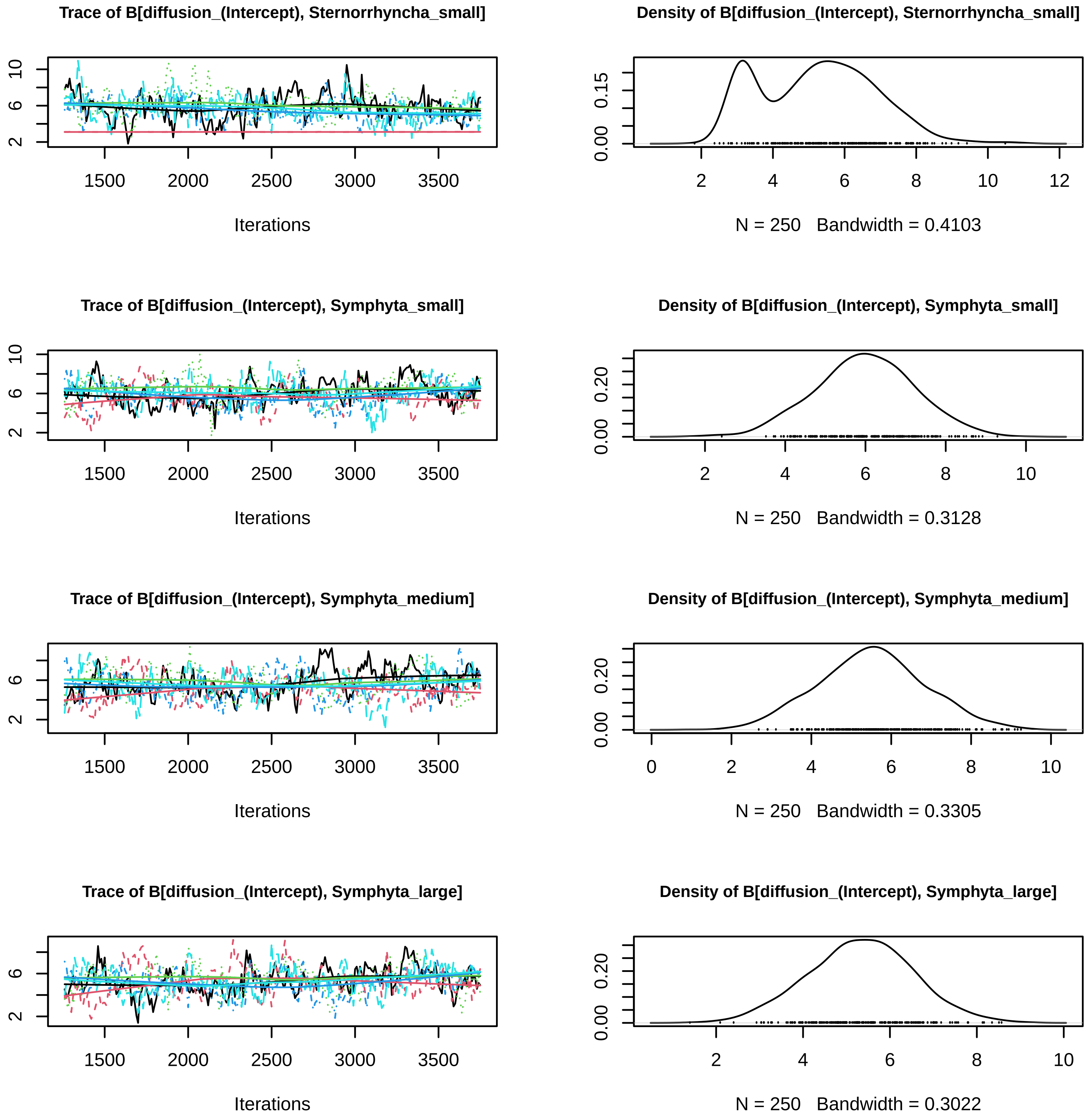
Trace of B[diffusion_(Intercept), Sternorrhyncha_small]
Density of B[diffusion_(Intercept), Sternorrhyncha_small]
Iterations
N = 250 Bandwidth = 0.4103
Trace of B[diffusion_(Intercept), Symphyta_small]
Density of B[diffusion_(Intercept), Symphyta_small]
Iterations
N = 250 Bandwidth = 0.3128
Trace of B[diffusion_(Intercept), Symphyta_medium]
Density of B[diffusion_(Intercept), Symphyta_medium]
Iterations
N = 250 Bandwidth = 0.3305
Trace of B[diffusion_(Intercept), Symphyta_large]
Density of B[diffusion_(Intercept), Symphyta_large]
Iterations
N = 250 Bandwidth = 0.3022

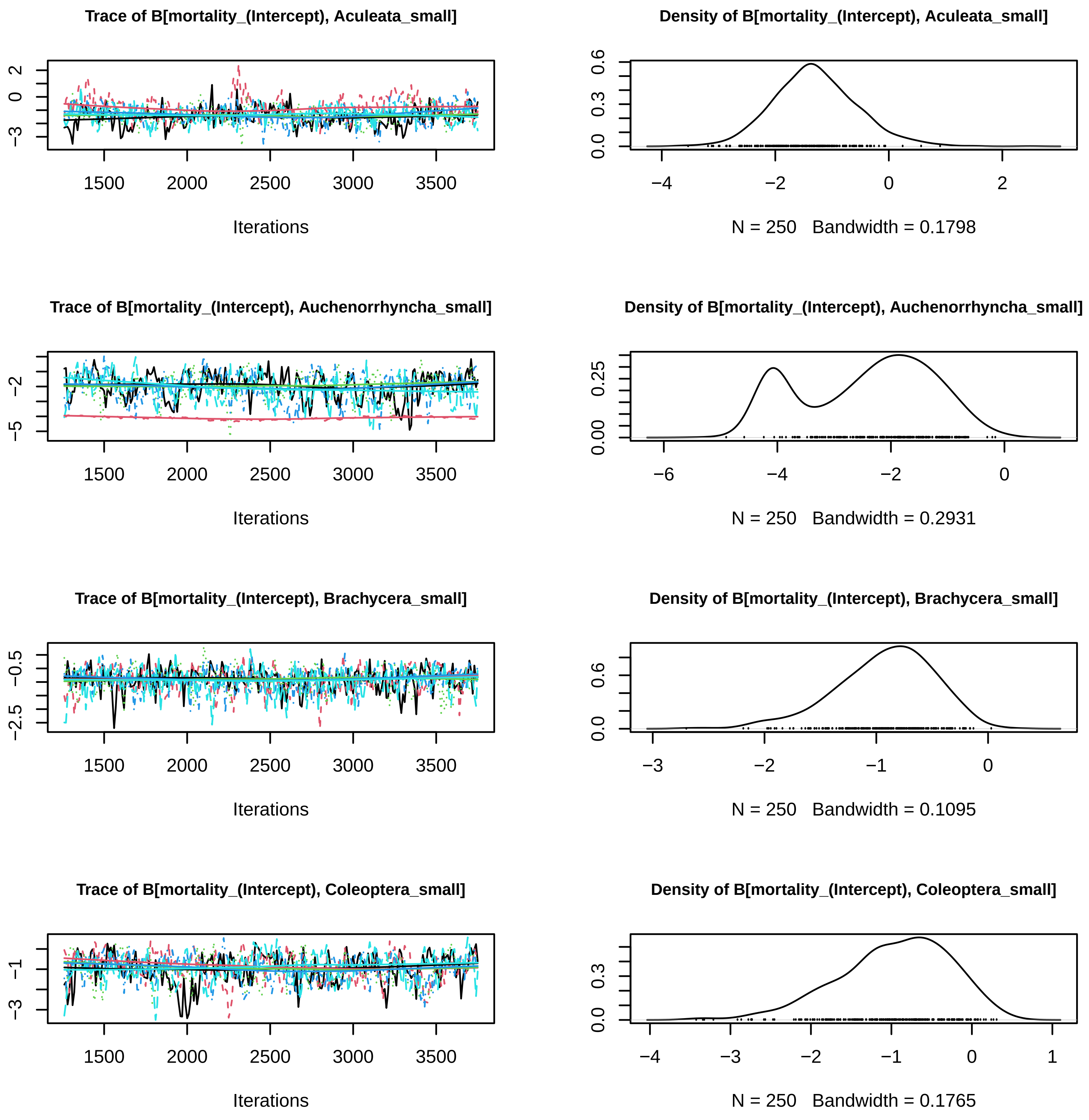
Trace of B[mortality_(Intercept), Aculeata_small]
Iterations
Density of B[mortality_(Intercept), Aculeata_small]
N = 250 Bandwidth = 0.1798
Trace of B[mortality_(Intercept), Auchenorrhyncha_small]
Iterations
Density of B[mortality_(Intercept), Auchenorrhyncha_small]
N = 250 Bandwidth = 0.2931
Trace of B[mortality_(Intercept), Brachycera_small]
Iterations
Density of B[mortality_(Intercept), Brachycera_small]
N = 250 Bandwidth = 0.1095
Trace of B[mortality_(Intercept), Coleoptera_small]
Iterations
Density of B[mortality_(Intercept), Coleoptera_small]
N = 250 Bandwidth = 0.1765

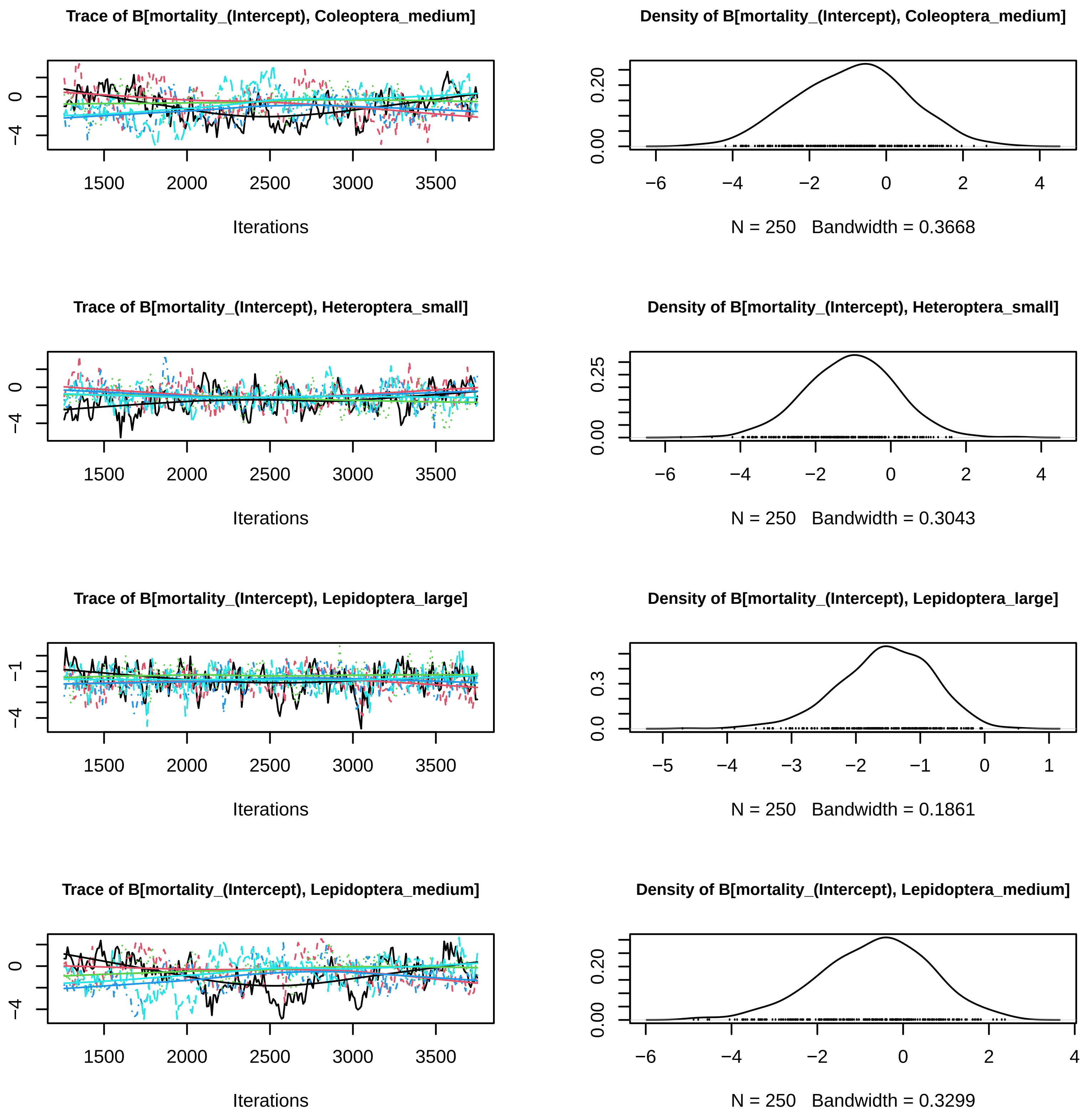
Trace of B[mortality_(Intercept), Coleoptera_medium]
Density of B[mortality_(Intercept), Coleoptera_medium]
Iterations
N = 250 Bandwidth = 0.3668
Trace of B[mortality_(Intercept), Heteroptera_small]
Density of B[mortality_(Intercept), Heteroptera_small]
Iterations
N = 250 Bandwidth = 0.3043
Trace of B[mortality_(Intercept), Lepidoptera_large]
Density of B[mortality_(Intercept), Lepidoptera_large]
Iterations
N = 250 Bandwidth = 0.1861
Trace of B[mortality_(Intercept), Lepidoptera_medium]
Density of B[mortality_(Intercept), Lepidoptera_medium]
Iterations
N = 250 Bandwidth = 0.3299

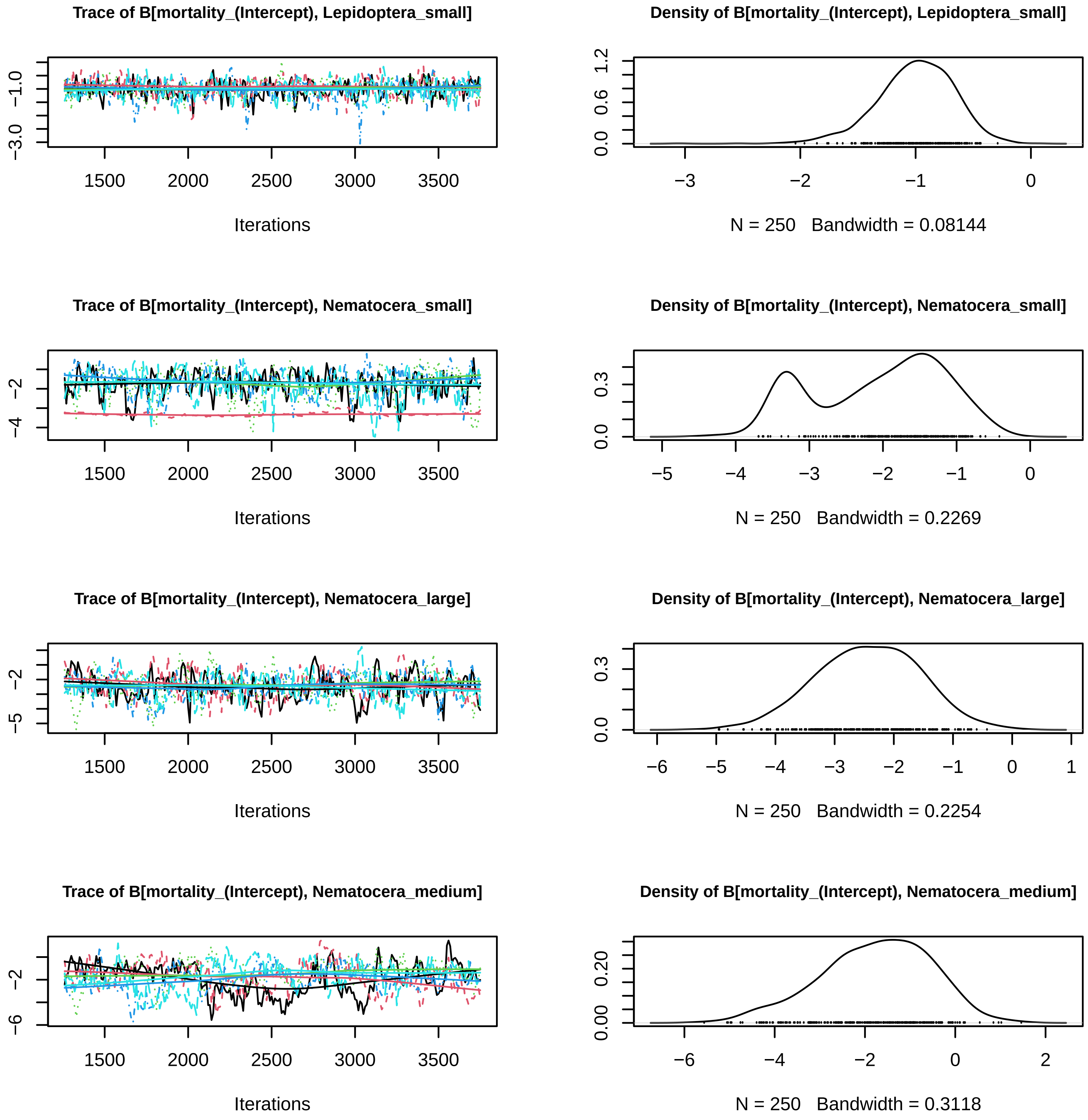

Trace of B[mortality_(Intercept), Lepidoptera_small]
Iterations
Density of B[mortality_(Intercept), Lepidoptera_small]
N = 250 Bandwidth = 0.08144
Trace of B[mortality_(Intercept), Nematocera_small]
Iterations
Density of B[mortality_(Intercept), Nematocera_small]
N = 250 Bandwidth = 0.2269
Trace of B[mortality_(Intercept), Nematocera_large]
Iterations
Density of B[mortality_(Intercept), Nematocera_large]
N = 250 Bandwidth = 0.2254
Trace of B[mortality_(Intercept), Nematocera_medium]
Iterations
Density of B[mortality_(Intercept), Nematocera_medium]
N = 250 Bandwidth = 0.3118

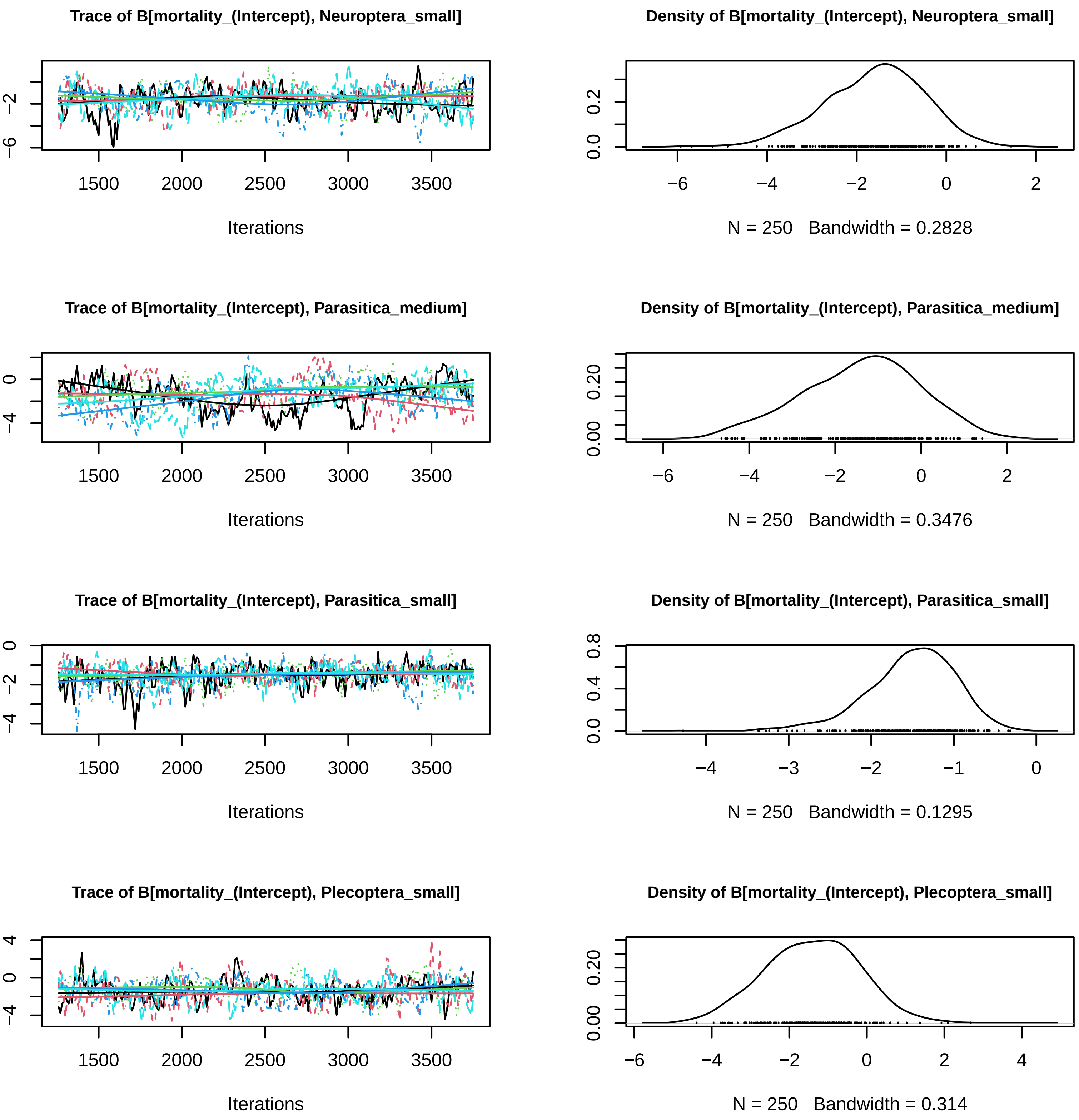
Trace of B[mortality_(Intercept), Neuroptera_small]
Density of B[mortality_(Intercept), Neuroptera_small]
Iterations
N = 250 Bandwidth = 0.2828
Trace of B[mortality_(Intercept), Parasitica_medium]
Density of B[mortality_(Intercept), Parasitica_medium]
Iterations
N = 250 Bandwidth = 0.3476
Trace of B[mortality_(Intercept), Parasitica_small]
Density of B[mortality_(Intercept), Parasitica_small]
Iterations
N = 250 Bandwidth = 0.1295
Trace of B[mortality_(Intercept), Plecoptera_small]
Density of B[mortality_(Intercept), Plecoptera_small]
Iterations
N = 250 Bandwidth = 0.314

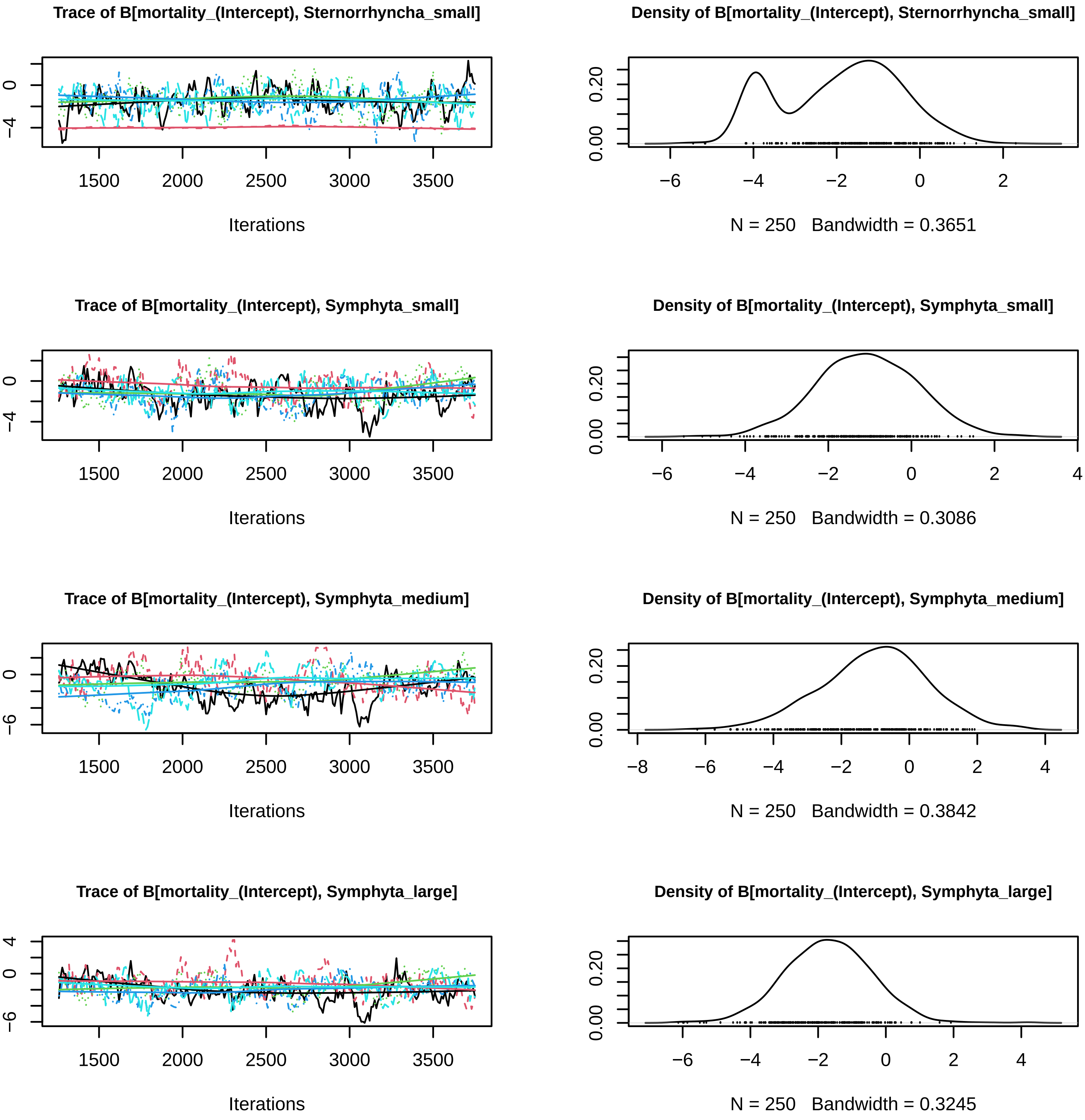
Trace of B[mortality_(Intercept), Sternorrhyncha_small]
Iterations
Density of B[mortality_(Intercept), Sternorrhyncha_small]
N = 250 Bandwidth = 0.3651
Trace of B[mortality_(Intercept), Symphyta_small]
Iterations
Density of B[mortality_(Intercept), Symphyta_small]
N = 250 Bandwidth = 0.3086
Trace of B[mortality_(Intercept), Symphyta_medium]
Iterations
Density of B[mortality_(Intercept), Symphyta_medium]
N = 250 Bandwidth = 0.3842
Trace of B[mortality_(Intercept), Symphyta_large]
Iterations
Density of B[mortality_(Intercept), Symphyta_large]
N = 250 Bandwidth = 0.3245

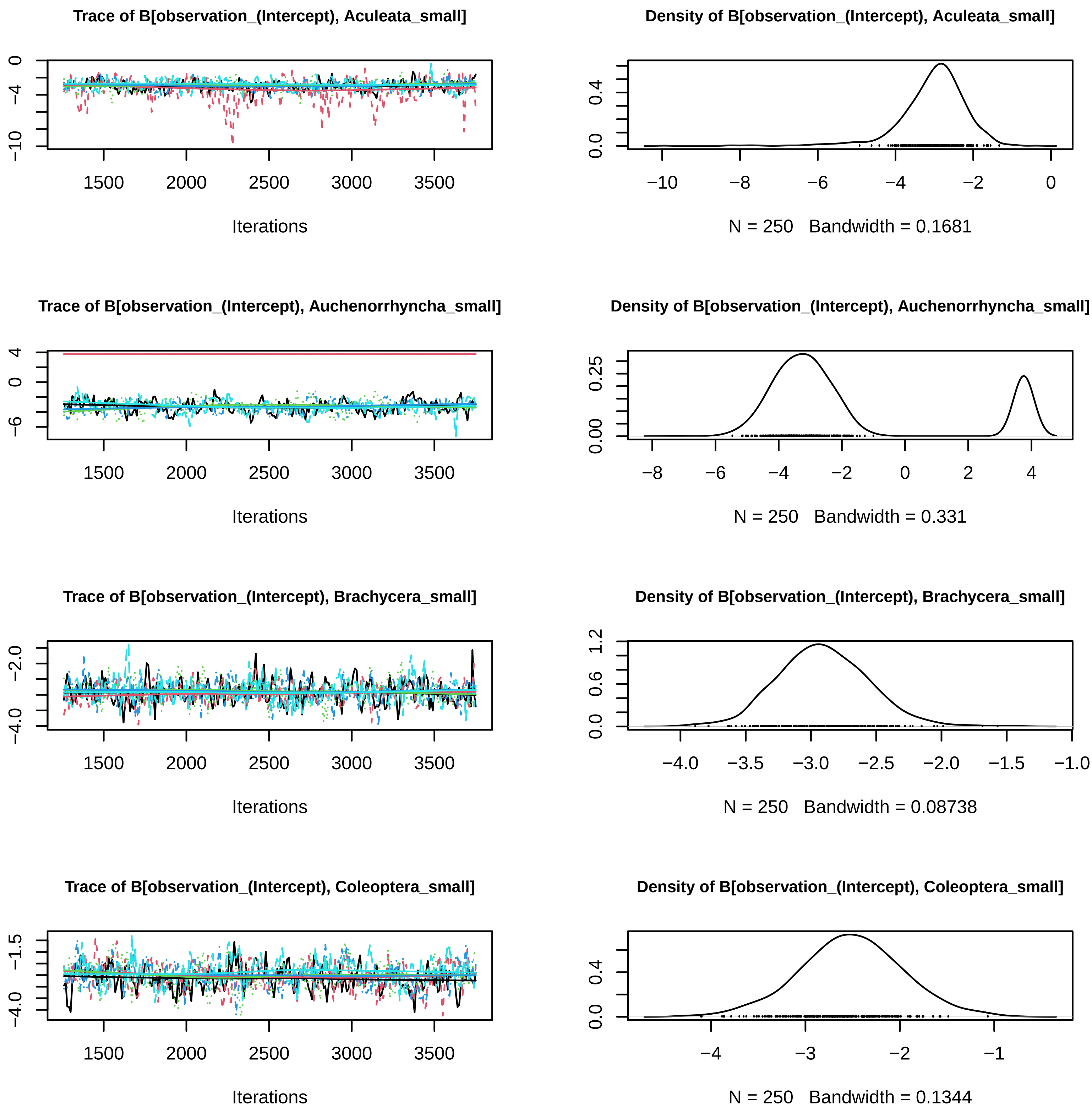

Trace of B[observation_(Intercept), Aculeata_small]
Iterations
Density of B[observation_(Intercept), Aculeata_small]
N = 250 Bandwidth = 0.1681
Trace of B[observation_(Intercept), Auchenorrhyncha_small]
Iterations
Density of B[observation_(Intercept), Auchenorrhyncha_small]
N = 250 Bandwidth = 0.331
Trace of B[observation_(Intercept), Brachycera_small]
Iterations
Density of B[observation_(Intercept), Brachycera_small]
N = 250 Bandwidth = 0.08738
Trace of B[observation_(Intercept), Coleoptera_small]
Iterations
Density of B[observation_(Intercept), Coleoptera_small]
N = 250 Bandwidth = 0.1344

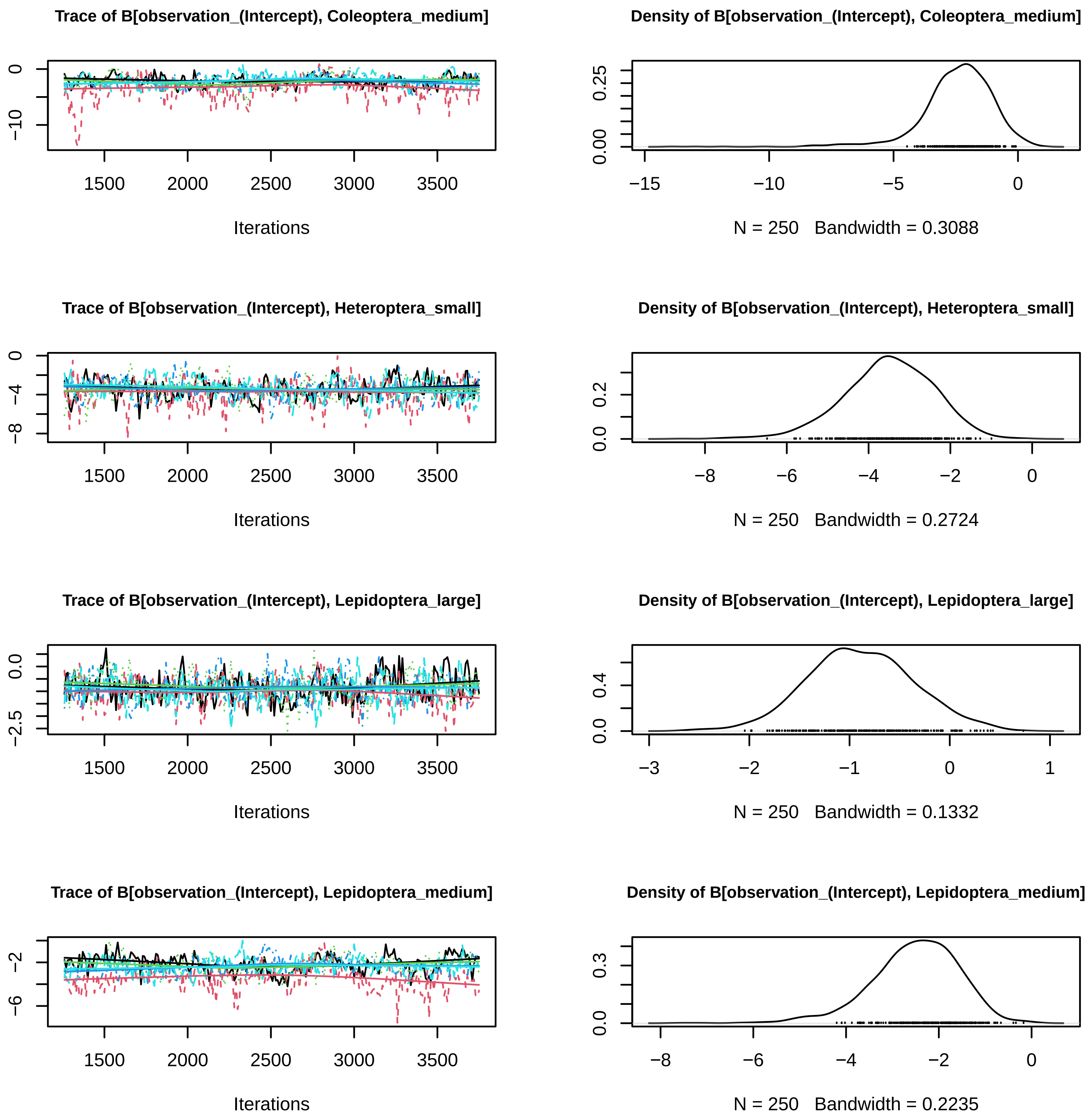
Trace of B[observation_(Intercept), Coleoptera_medium]
Iterations
Density of B[observation_(Intercept), Coleoptera_medium]
N = 250 Bandwidth = 0.3088
Trace of B[observation_(Intercept), Heteroptera_small]
Iterations
Density of B[observation_(Intercept), Heteroptera_small]
N = 250 Bandwidth = 0.2724
Trace of B[observation_(Intercept), Lepidoptera_large]
Iterations
Density of B[observation_(Intercept), Lepidoptera_large]
N = 250 Bandwidth = 0.1332
Trace of B[observation_(Intercept), Lepidoptera_medium]
Iterations
Density of B[observation_(Intercept), Lepidoptera_medium]
N = 250 Bandwidth = 0.2235

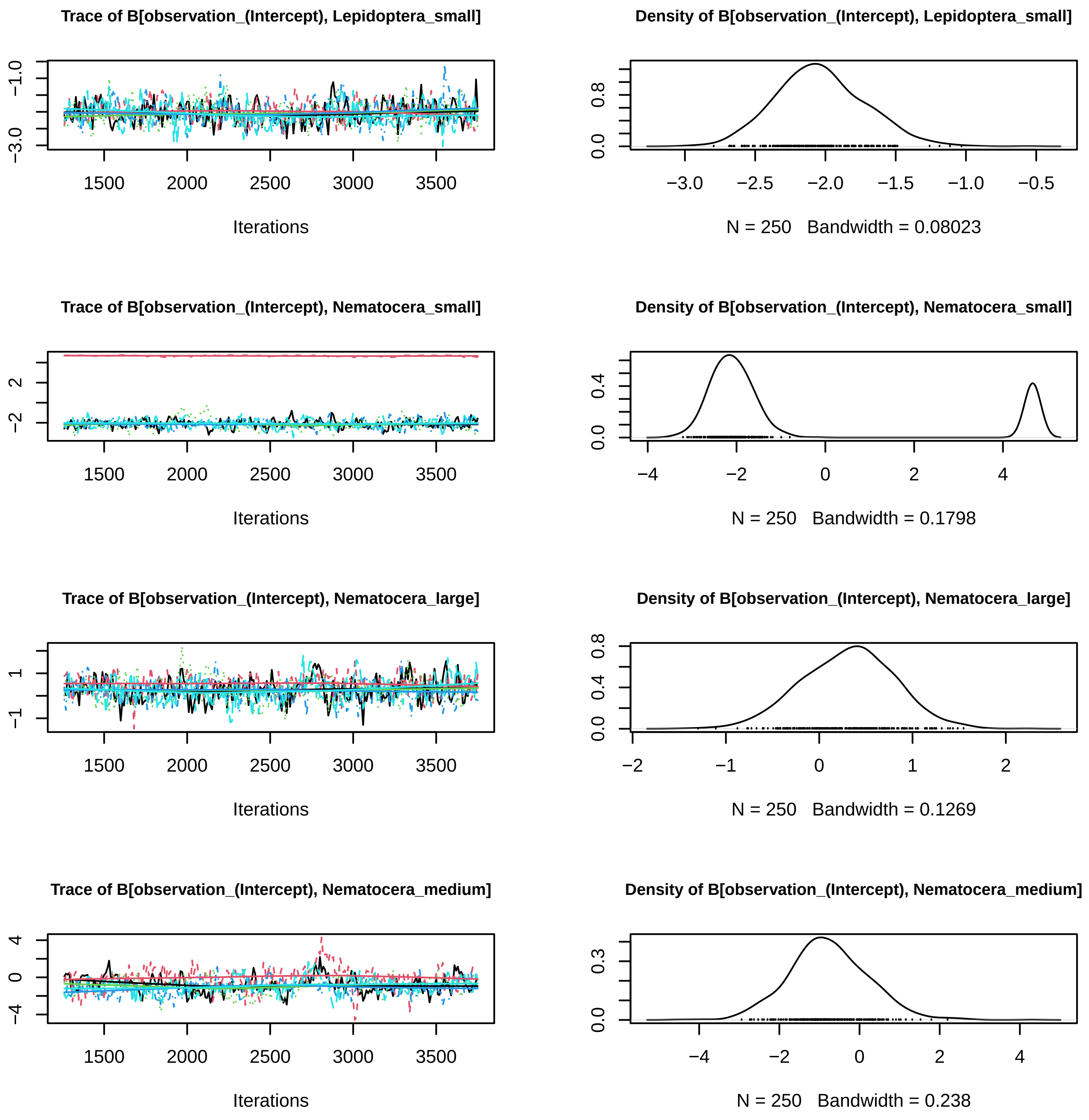
Trace of B[observation_(Intercept), Lepidoptera_small]
Density of B[observation_(Intercept), Lepidoptera_small]
−3.0
−1.0
1500
2000
2500
3000
3500
Iterations
0.0
0.8
−3.0
−2.5
−2.0
−1.5
−1.0
−0.5
N = 250 Bandwidth = 0.08023
Trace of B[observation_(Intercept), Nematocera_small]
Density of B[observation_(Intercept), Nematocera_small]
−2
2
1500
2000
2500
3000
3500
Iterations
0.0
0.4
−4
−2
0
2
4
N = 250 Bandwidth = 0.1798
Trace of B[observation_(Intercept), Nematocera_large]
Density of B[observation_(Intercept), Nematocera_large]
−1
1
1500
2000
2500
3000
3500
Iterations
0.0
0.4
0.8
−2
−1
0
1
2
N = 250 Bandwidth = 0.1269
Trace of B[observation_(Intercept), Nematocera_medium]
Density of B[observation_(Intercept), Nematocera_medium]
−4
0
4
1500
2000
2500
3000
3500
Iterations
0.0
0.3
−4
−2
0
2
4
N = 250 Bandwidth = 0.238

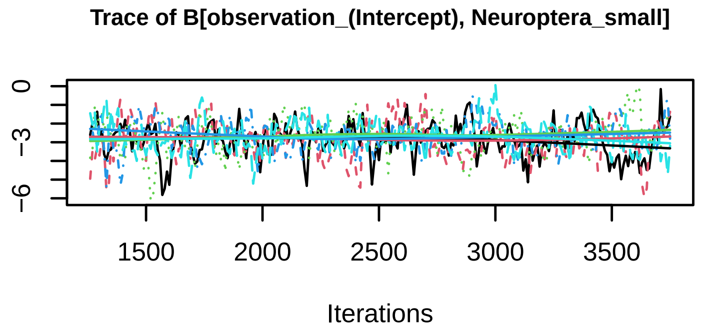
Trace of B[observation_(Intercept), Neuroptera_small]
0
−3
−6
1500
2000
2500
3000
3500
Iterations

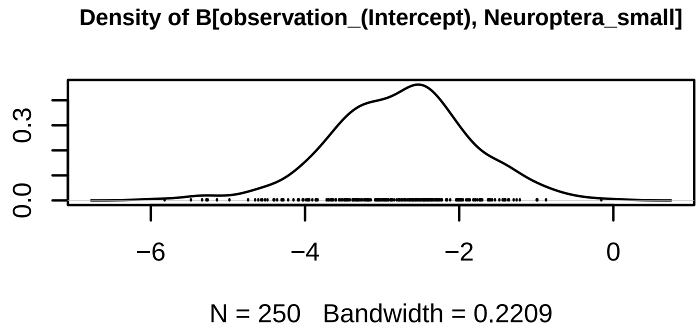
Density of B[observation_(Intercept), Neuroptera_small]
0.0
0.3
−6
−4
−2
0
N = 250 Bandwidth = 0.2209

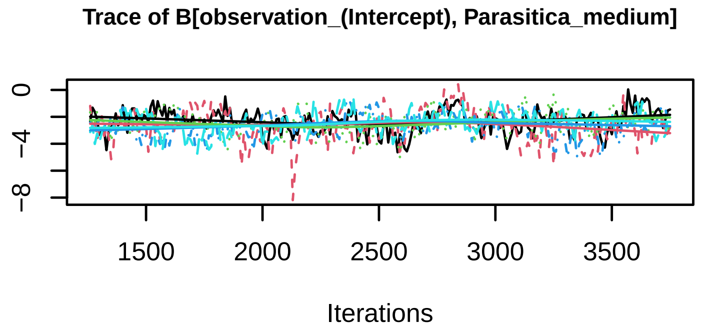
Trace of B[observation_(Intercept), Parasitica_medium]
0
−4
−8
1500
2000
2500
3000
3500
Iterations

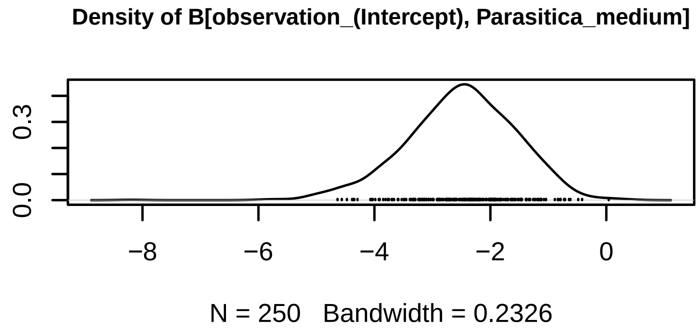
Density of B[observation_(Intercept), Parasitica_medium]
0.0
0.3
−8
−6
−4
−2
0
N = 250 Bandwidth = 0.2326

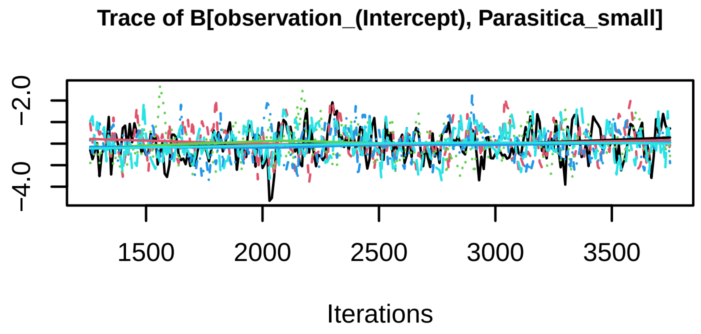
Trace of B[observation_(Intercept), Parasitica_small]
−2.0
−4.0
1500
2000
2500
3000
3500
Iterations

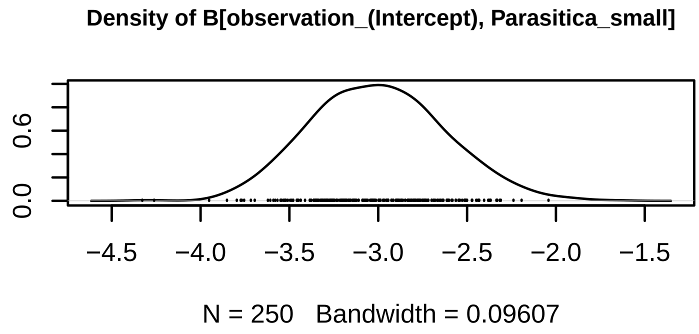
Density of B[observation_(Intercept), Parasitica_small]
0.0
0.6
−4.5
−4.0
−3.5
−3.0
−2.5
−2.0
−1.5
N = 250 Bandwidth = 0.09607

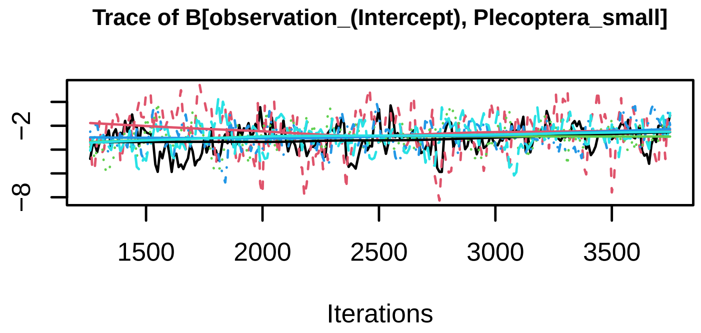
Trace of B[observation_(Intercept), Plecoptera_small]
−2
−8
1500
2000
2500
3000
3500
Iterations

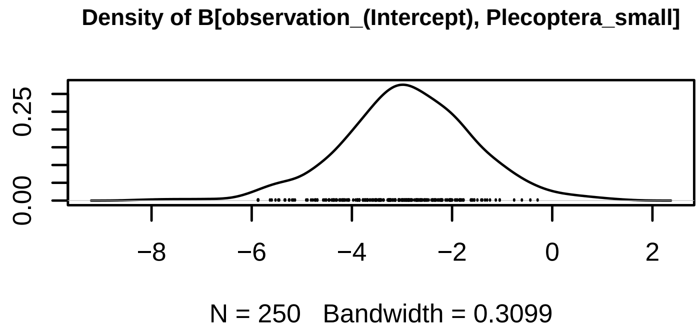
Density of B[observation_(Intercept), Plecoptera_small]
0.00
0.25
−8
−6
−4
−2
0
2
N = 250 Bandwidth = 0.3099

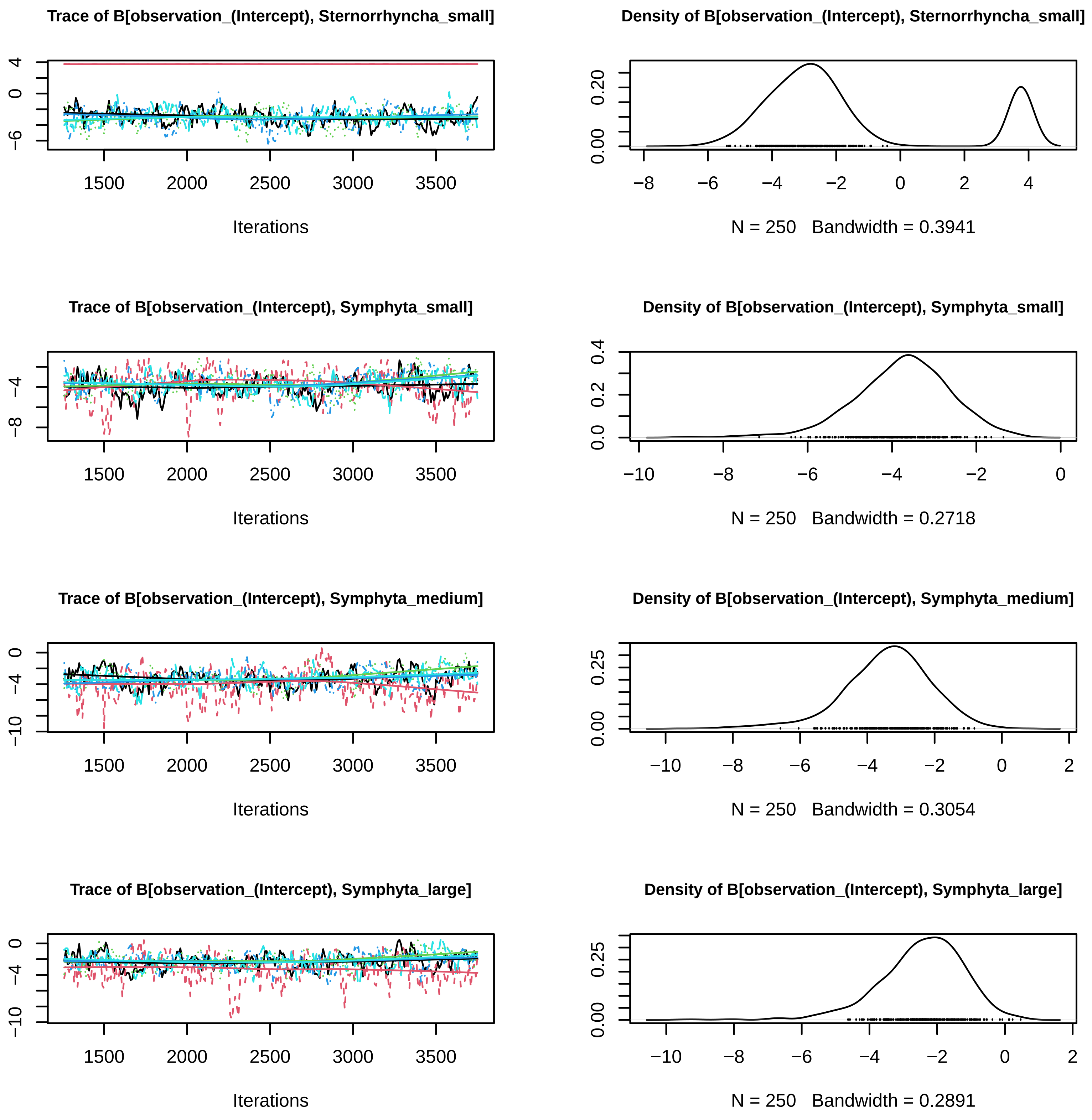

Trace of B[observation_(Intercept), Sternorrhyncha_small]
Iterations
Density of B[observation_(Intercept), Sternorrhyncha_small]
N = 250 Bandwidth = 0.3941
Trace of B[observation_(Intercept), Symphyta_small]
Iterations
Density of B[observation_(Intercept), Symphyta_small]
N = 250 Bandwidth = 0.2718
Trace of B[observation_(Intercept), Symphyta_medium]
Iterations
Density of B[observation_(Intercept), Symphyta_medium]
N = 250 Bandwidth = 0.3054
Trace of B[observation_(Intercept), Symphyta_large]
Iterations
Density of B[observation_(Intercept), Symphyta_large]
N = 250 Bandwidth = 0.2891

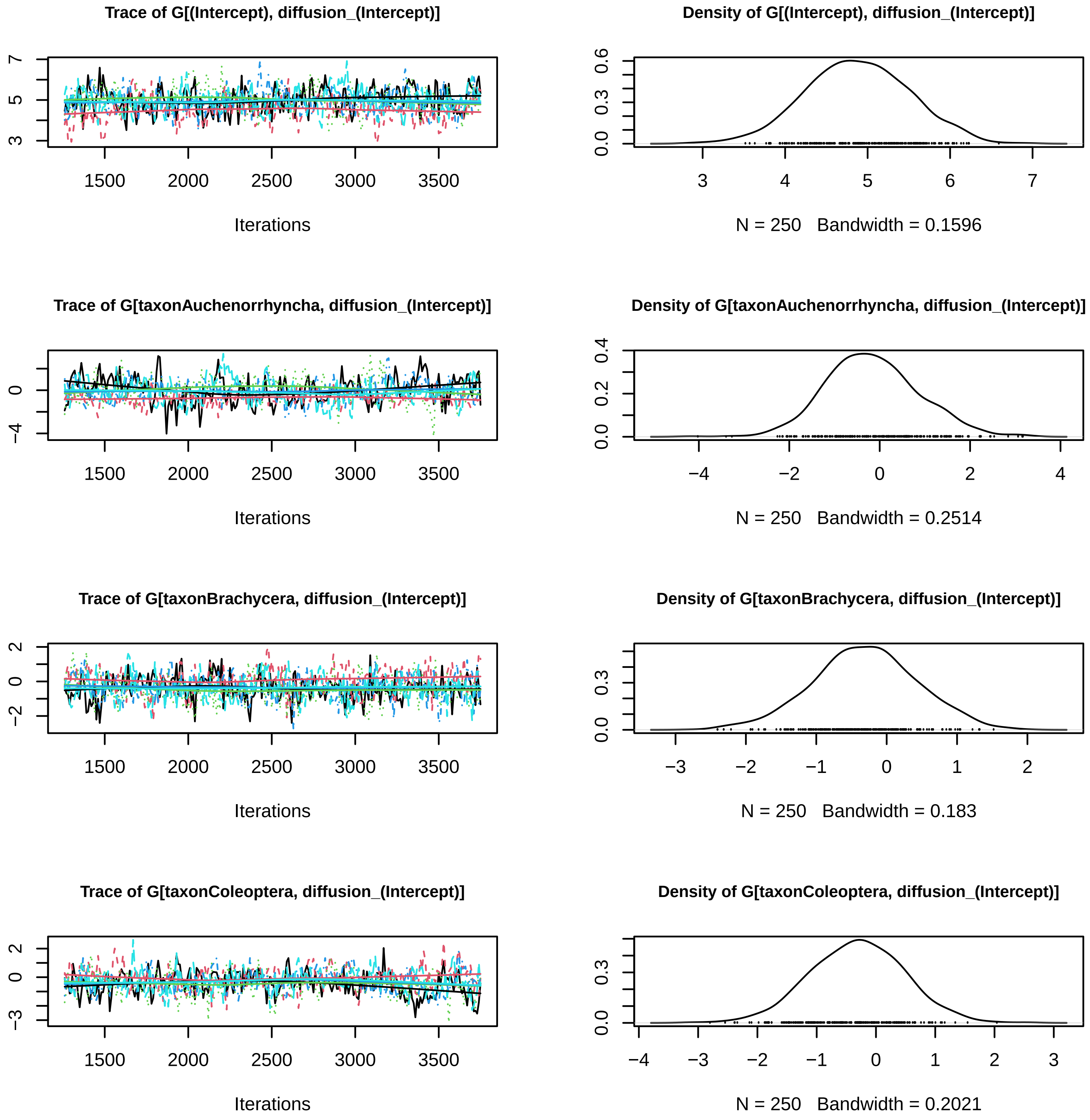

Trace of G[(Intercept), diffusion_(Intercept)]
Iterations
Density of G[(Intercept), diffusion_(Intercept)]
N = 250 Bandwidth = 0.1596
Trace of G[taxonAuchenorrhyncha, diffusion_(Intercept)]
Iterations
Density of G[taxonAuchenorrhyncha, diffusion_(Intercept)]
N = 250 Bandwidth = 0.2514
Trace of G[taxonBrachycera, diffusion_(Intercept)]
Iterations
Density of G[taxonBrachycera, diffusion_(Intercept)]
N = 250 Bandwidth = 0.183
Trace of G[taxonColeoptera, diffusion_(Intercept)]
Iterations
Density of G[taxonColeoptera, diffusion_(Intercept)]
N = 250 Bandwidth = 0.2021

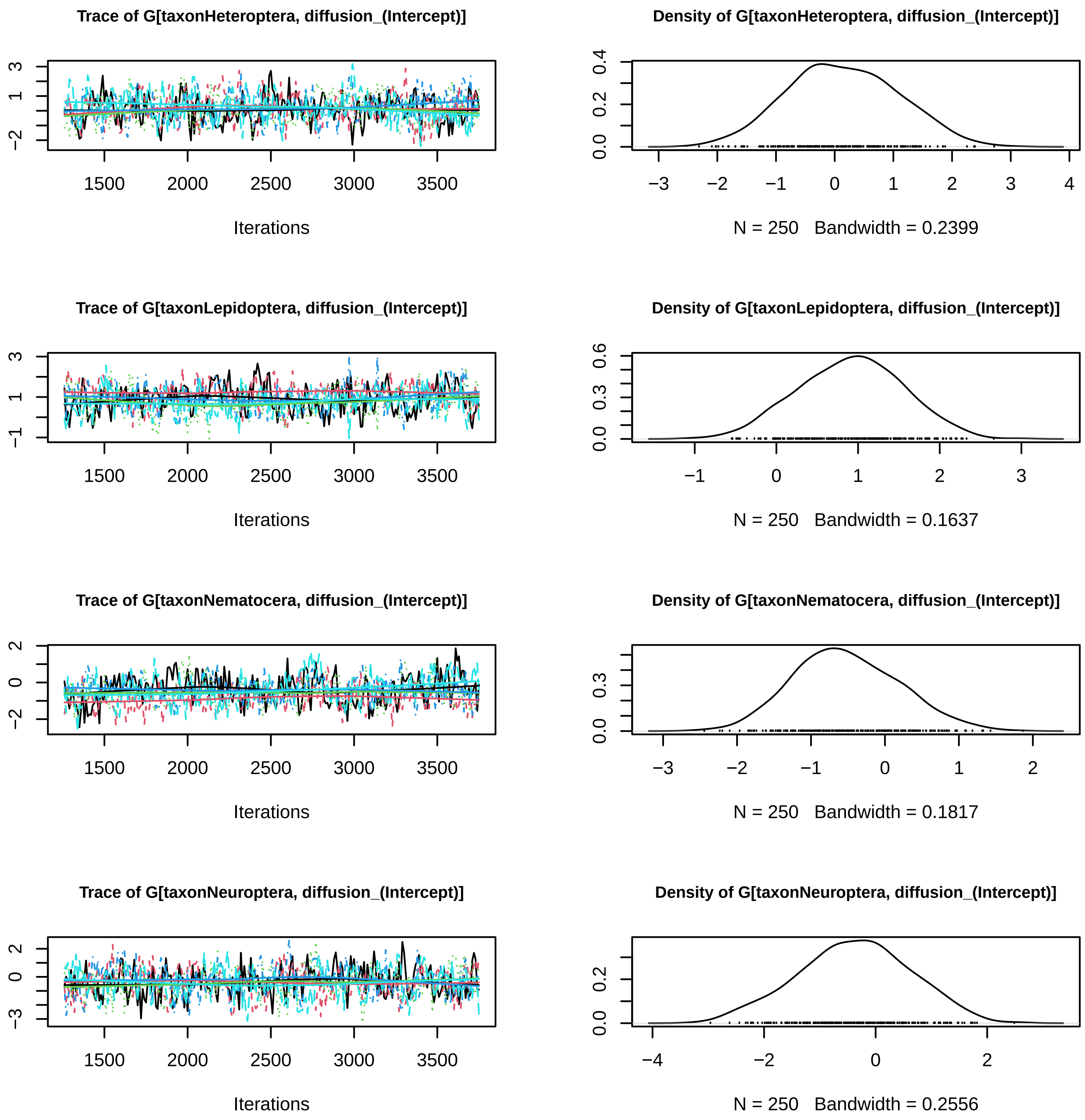

Trace of G[taxonHeteroptera, diffusion_(Intercept)]
Iterations
Density of G[taxonHeteroptera, diffusion_(Intercept)]
N = 250 Bandwidth = 0.2399
Trace of G[taxonLepidoptera, diffusion_(Intercept)]
Iterations
Density of G[taxonLepidoptera, diffusion_(Intercept)]
N = 250 Bandwidth = 0.1637
Trace of G[taxonNematocera, diffusion_(Intercept)]
Iterations
Density of G[taxonNematocera, diffusion_(Intercept)]
N = 250 Bandwidth = 0.1817
Trace of G[taxonNeuroptera, diffusion_(Intercept)]
Iterations
Density of G[taxonNeuroptera, diffusion_(Intercept)]
N = 250 Bandwidth = 0.2556

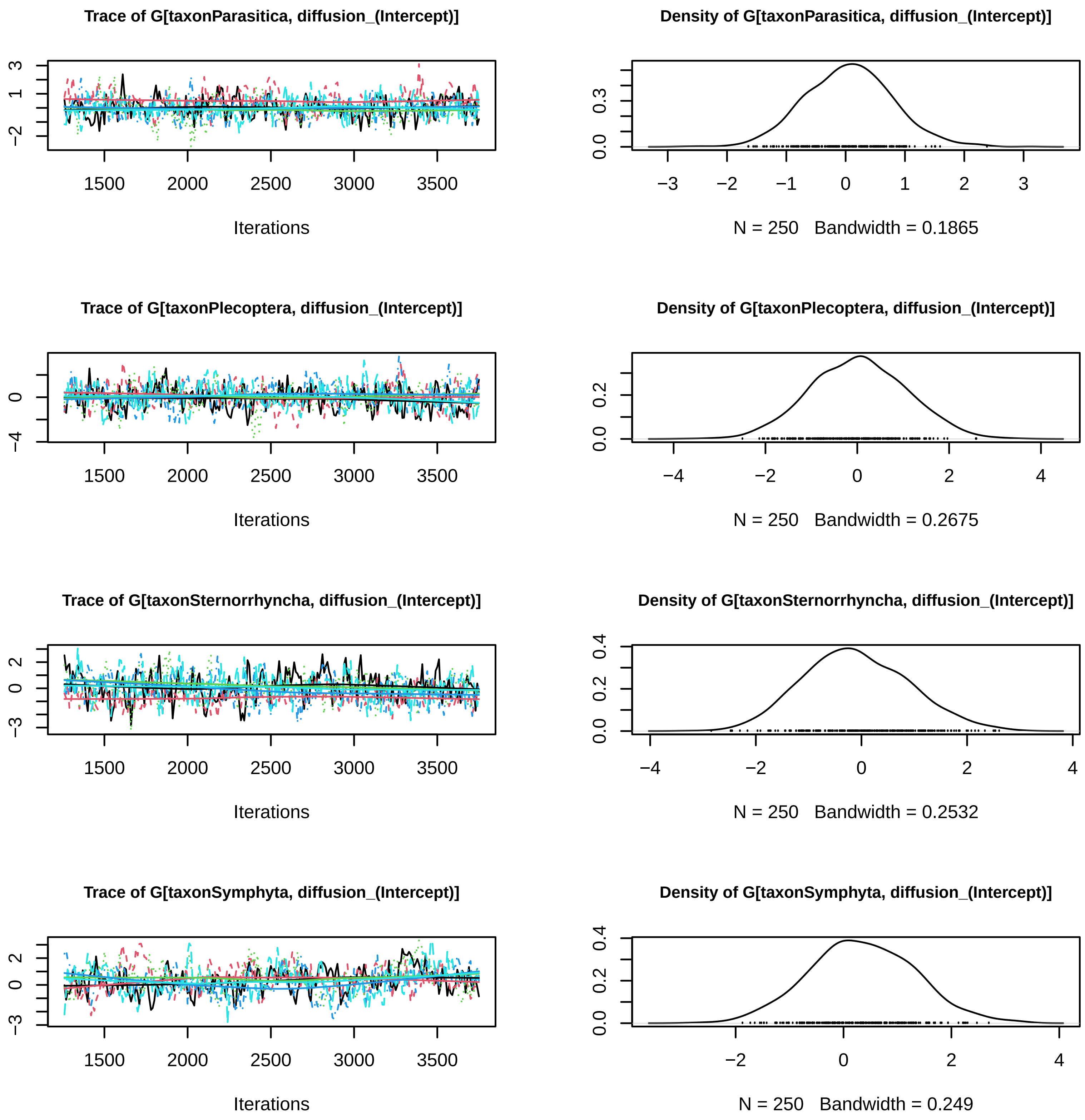
Trace of G[taxonParasitica, diffusion_(Intercept)]
Iterations
Density of G[taxonParasitica, diffusion_(Intercept)]
N = 250 Bandwidth = 0.1865
Trace of G[taxonPlecoptera, diffusion_(Intercept)]
Iterations
Density of G[taxonPlecoptera, diffusion_(Intercept)]
N = 250 Bandwidth = 0.2675
Trace of G[taxonSternorrhyncha, diffusion_(Intercept)]
Iterations
Density of G[taxonSternorrhyncha, diffusion_(Intercept)]
N = 250 Bandwidth = 0.2532
Trace of G[taxonSymphyta, diffusion_(Intercept)]
Iterations
Density of G[taxonSymphyta, diffusion_(Intercept)]
N = 250 Bandwidth = 0.249

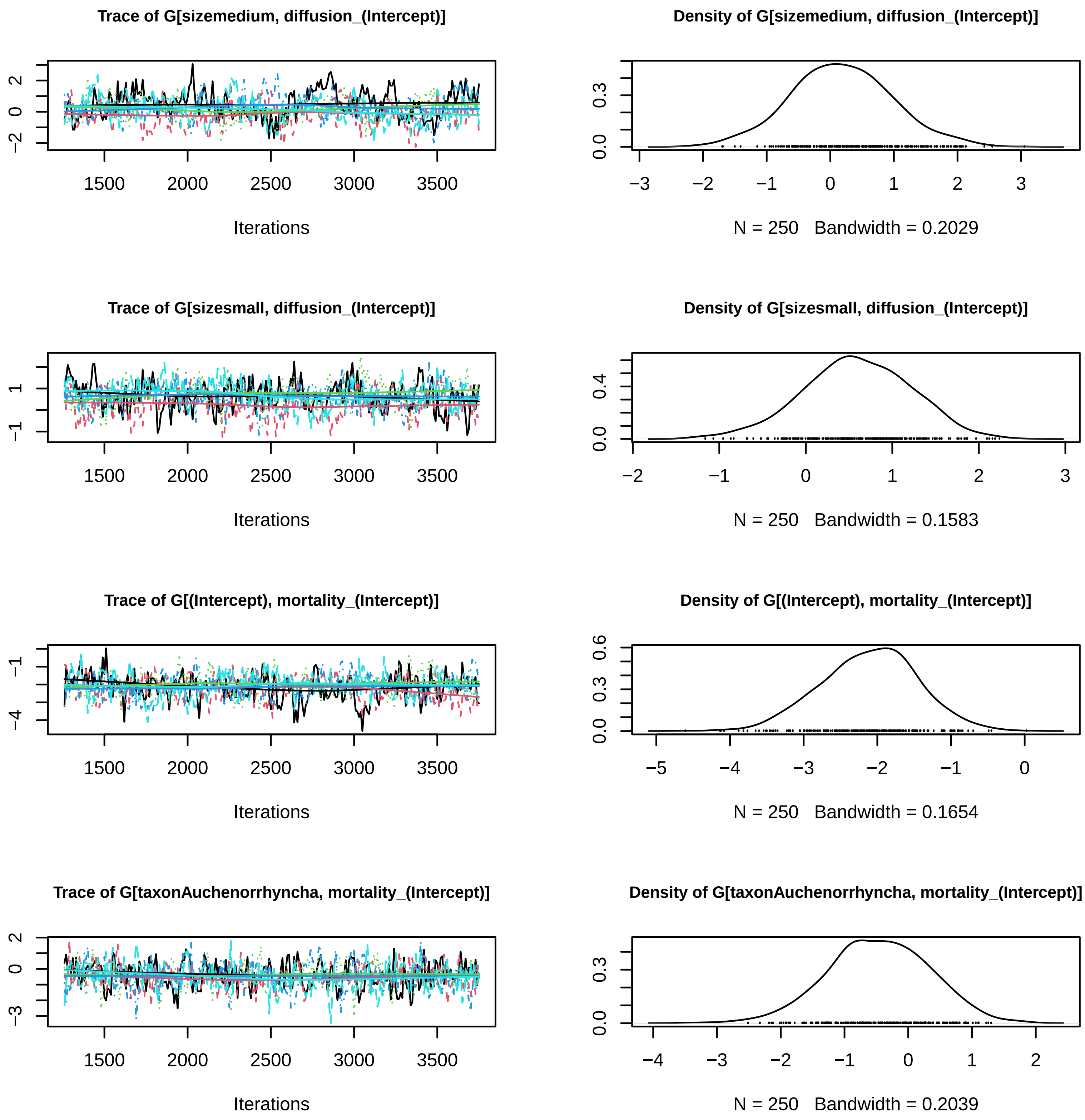
Trace of G[sizemedium, diffusion_(Intercept)]
Iterations
Density of G[sizemedium, diffusion_(Intercept)]
N = 250   Bandwidth = 0.2029
Trace of G[sizesmall, diffusion_(Intercept)]
Iterations
Density of G[sizesmall, diffusion_(Intercept)]
N = 250   Bandwidth = 0.1583
Trace of G[(Intercept), mortality_(Intercept)]
Iterations
Density of G[(Intercept), mortality_(Intercept)]
N = 250   Bandwidth = 0.1654
Trace of G[taxonAuchenorrhyncha, mortality_(Intercept)]
Iterations
Density of G[taxonAuchenorrhyncha, mortality_(Intercept)]
N = 250   Bandwidth = 0.2039

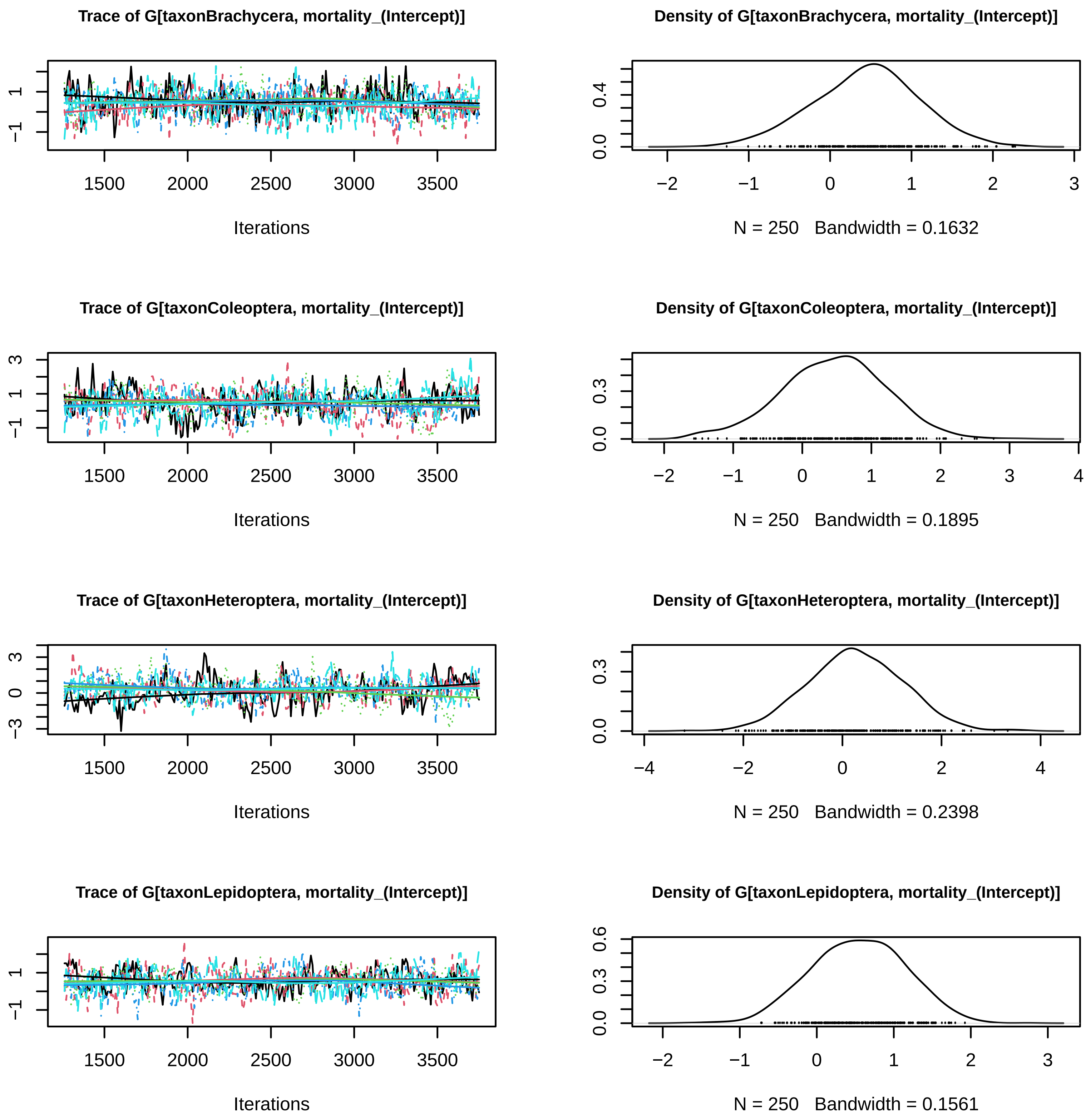
Trace of G[taxonBrachycera, mortality_(Intercept)]
Iterations
Density of G[taxonBrachycera, mortality_(Intercept)]
N = 250 Bandwidth = 0.1632
Trace of G[taxonColeoptera, mortality_(Intercept)]
Iterations
Density of G[taxonColeoptera, mortality_(Intercept)]
N = 250 Bandwidth = 0.1895
Trace of G[taxonHeteroptera, mortality_(Intercept)]
Iterations
Density of G[taxonHeteroptera, mortality_(Intercept)]
N = 250 Bandwidth = 0.2398
Trace of G[taxonLepidoptera, mortality_(Intercept)]
Iterations
Density of G[taxonLepidoptera, mortality_(Intercept)]
N = 250 Bandwidth = 0.1561

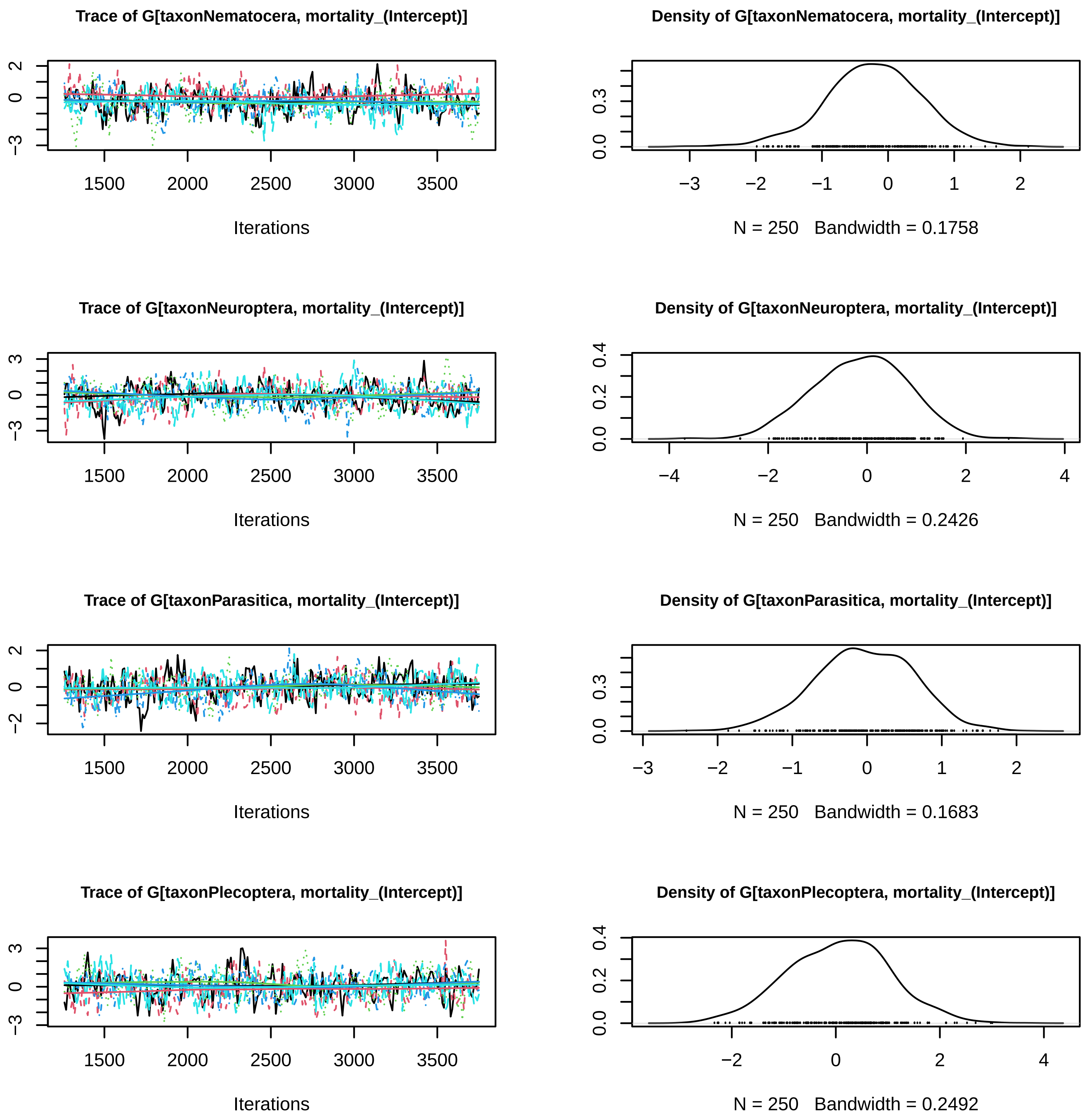

Trace of G[taxonNematocera, mortality_(Intercept)]
Iterations
Density of G[taxonNematocera, mortality_(Intercept)]
N = 250 Bandwidth = 0.1758
Trace of G[taxonNeuroptera, mortality_(Intercept)]
Iterations
Density of G[taxonNeuroptera, mortality_(Intercept)]
N = 250 Bandwidth = 0.2426
Trace of G[taxonParasitica, mortality_(Intercept)]
Iterations
Density of G[taxonParasitica, mortality_(Intercept)]
N = 250 Bandwidth = 0.1683
Trace of G[taxonPlecoptera, mortality_(Intercept)]
Iterations
Density of G[taxonPlecoptera, mortality_(Intercept)]
N = 250 Bandwidth = 0.2492

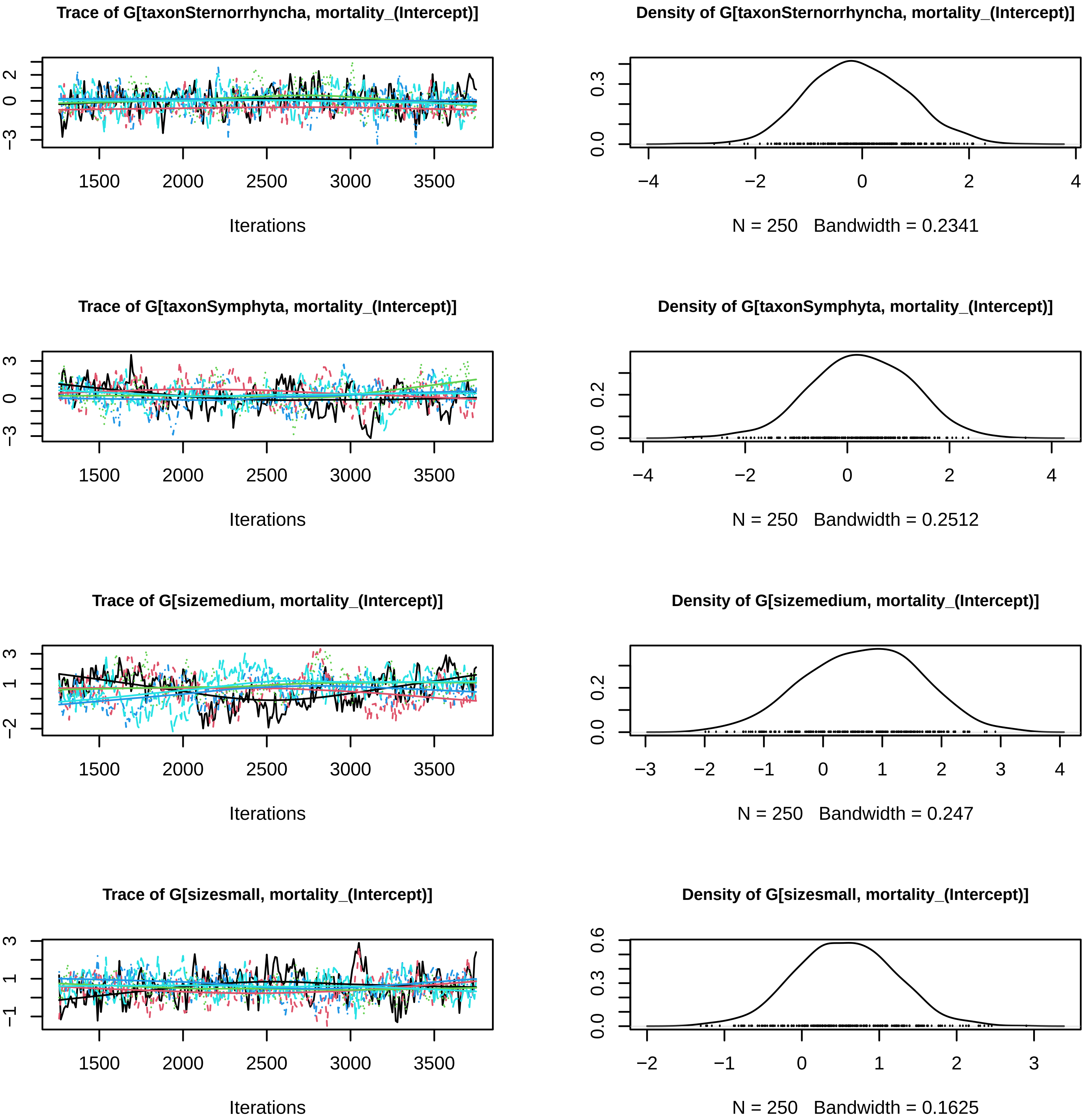
Trace of G[taxonSternorrhyncha, mortality_(Intercept)]
Iterations
Density of G[taxonSternorrhyncha, mortality_(Intercept)]
N = 250 Bandwidth = 0.2341
Trace of G[taxonSymphyta, mortality_(Intercept)]
Iterations
Density of G[taxonSymphyta, mortality_(Intercept)]
N = 250 Bandwidth = 0.2512
Trace of G[sizemedium, mortality_(Intercept)]
Iterations
Density of G[sizemedium, mortality_(Intercept)]
N = 250 Bandwidth = 0.247
Trace of G[sizesmall, mortality_(Intercept)]
Iterations
Density of G[sizesmall, mortality_(Intercept)]
N = 250 Bandwidth = 0.1625

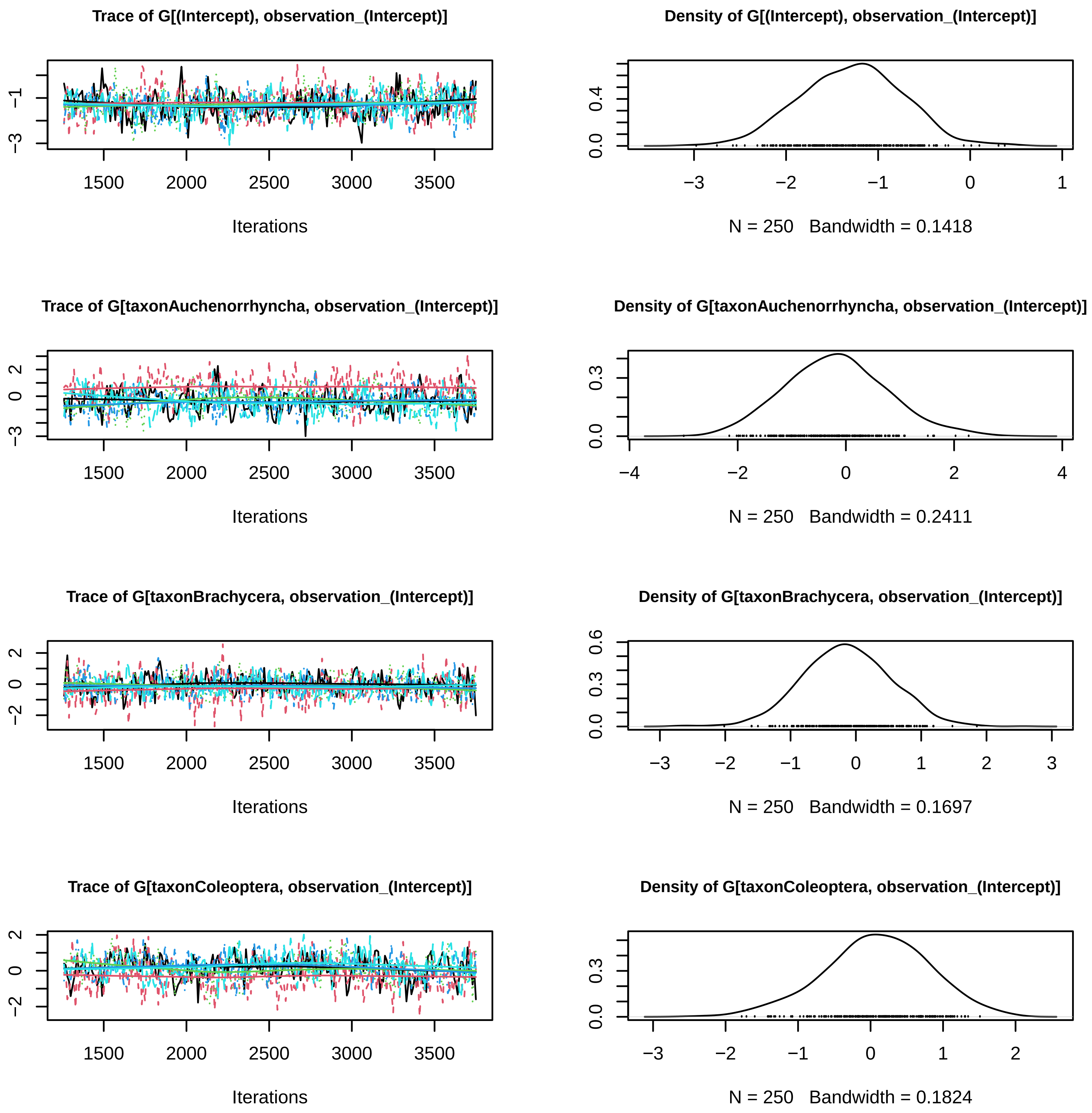
Trace of G[(Intercept), observation_(Intercept)]
Iterations
Density of G[(Intercept), observation_(Intercept)]
N = 250 Bandwidth = 0.1418
Trace of G[taxonAuchenorrhyncha, observation_(Intercept)]
Iterations
Density of G[taxonAuchenorrhyncha, observation_(Intercept)]
N = 250 Bandwidth = 0.2411
Trace of G[taxonBrachycera, observation_(Intercept)]
Iterations
Density of G[taxonBrachycera, observation_(Intercept)]
N = 250 Bandwidth = 0.1697
Trace of G[taxonColeoptera, observation_(Intercept)]
Iterations
Density of G[taxonColeoptera, observation_(Intercept)]
N = 250 Bandwidth = 0.1824

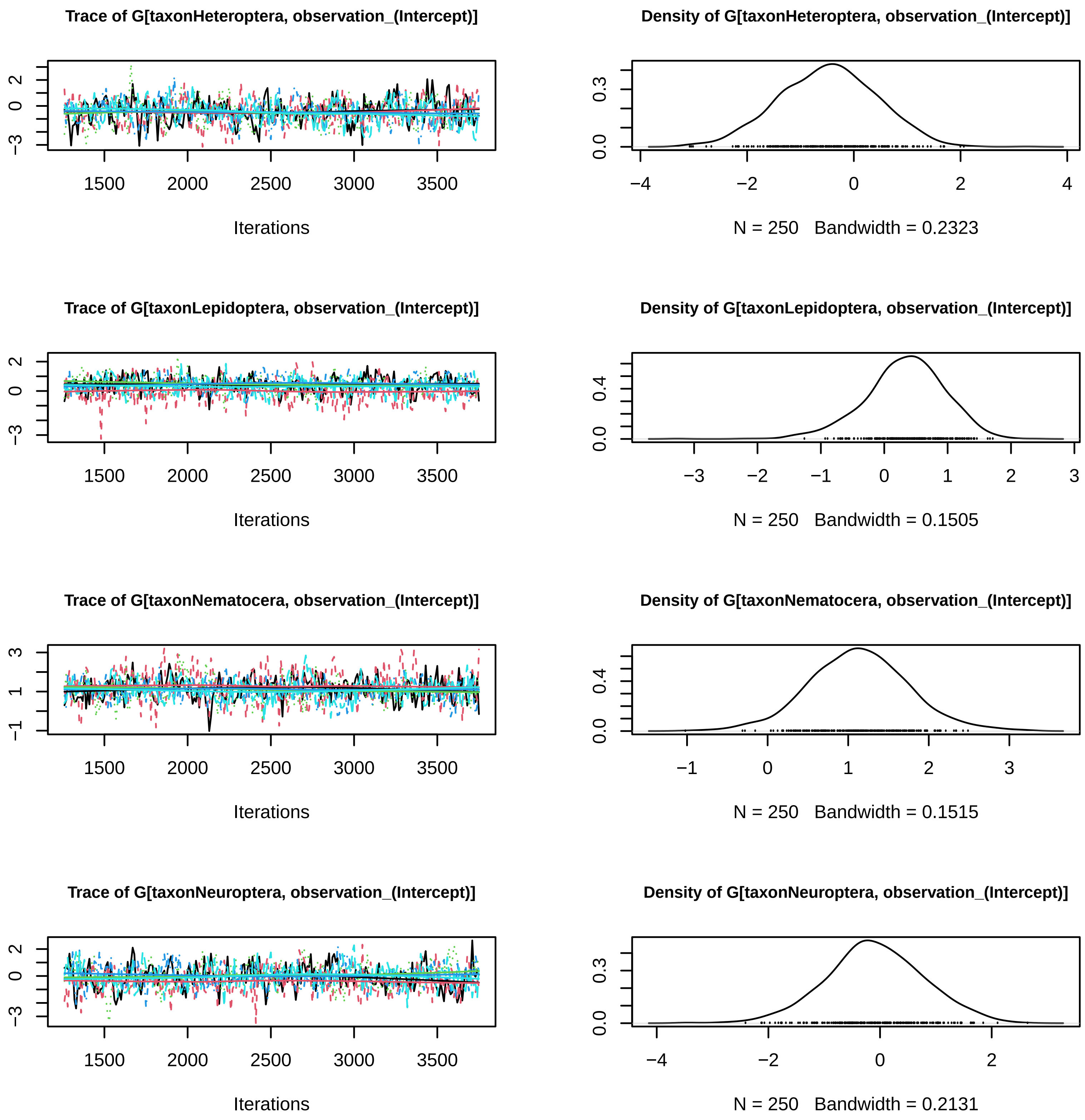
Trace of G[taxonHeteroptera, observation_(Intercept)]
Iterations
Density of G[taxonHeteroptera, observation_(Intercept)]
N = 250 Bandwidth = 0.2323
Trace of G[taxonLepidoptera, observation_(Intercept)]
Iterations
Density of G[taxonLepidoptera, observation_(Intercept)]
N = 250 Bandwidth = 0.1505
Trace of G[taxonNematocera, observation_(Intercept)]
Iterations
Density of G[taxonNematocera, observation_(Intercept)]
N = 250 Bandwidth = 0.1515
Trace of G[taxonNeuroptera, observation_(Intercept)]
Iterations
Density of G[taxonNeuroptera, observation_(Intercept)]
N = 250 Bandwidth = 0.2131

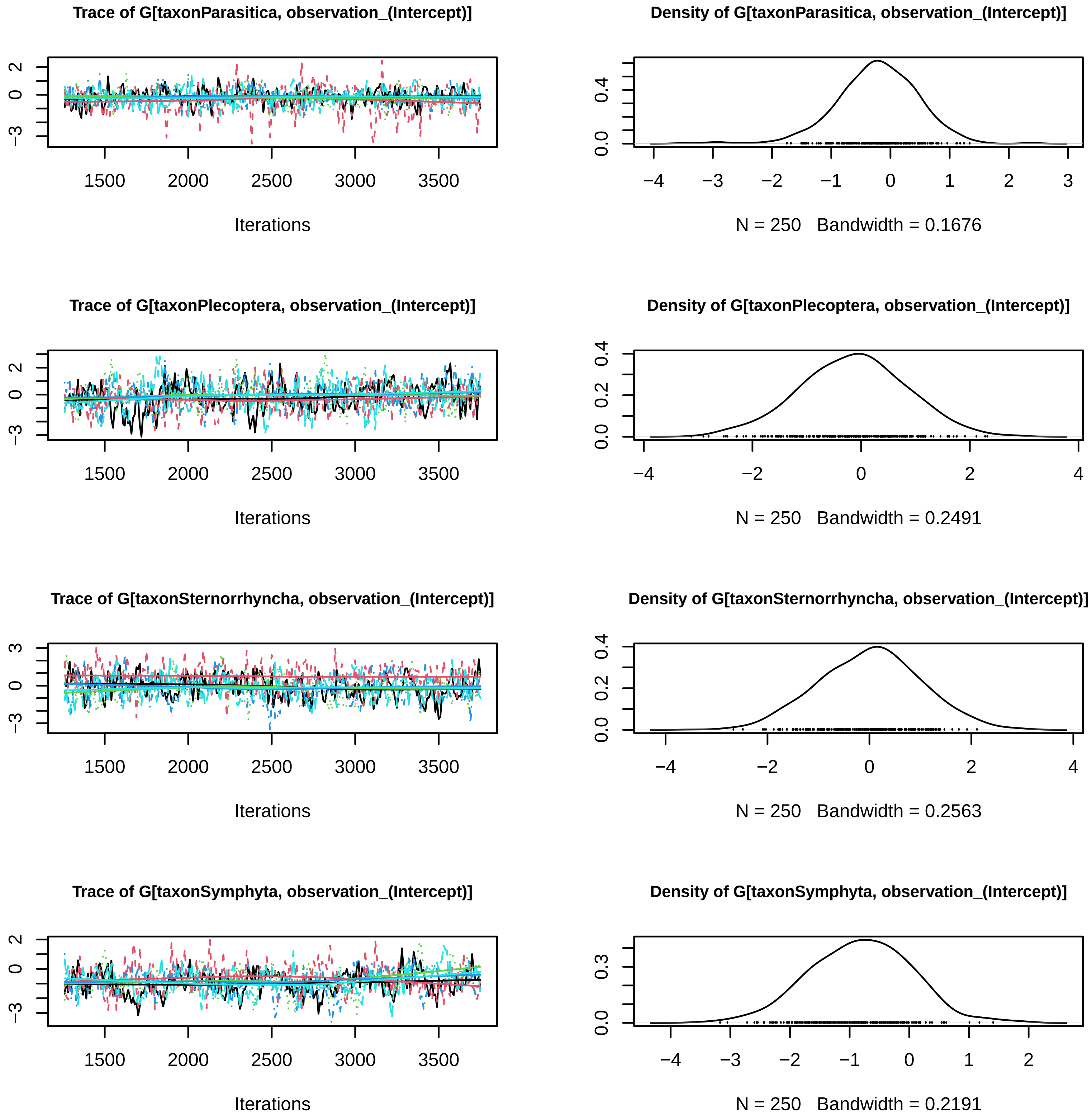

Trace of G[taxonParasitica, observation_(Intercept)]
Iterations
N = 250 Bandwidth = 0.1676
Density of G[taxonParasitica, observation_(Intercept)]
Trace of G[taxonPlecoptera, observation_(Intercept)]
Iterations
Density of G[taxonPlecoptera, observation_(Intercept)]
N = 250 Bandwidth = 0.2491
Trace of G[taxonSternorrhyncha, observation_(Intercept)]
Iterations
Density of G[taxonSternorrhyncha, observation_(Intercept)]
N = 250 Bandwidth = 0.2563
Trace of G[taxonSymphyta, observation_(Intercept)]
Iterations
Density of G[taxonSymphyta, observation_(Intercept)]
N = 250 Bandwidth = 0.2191

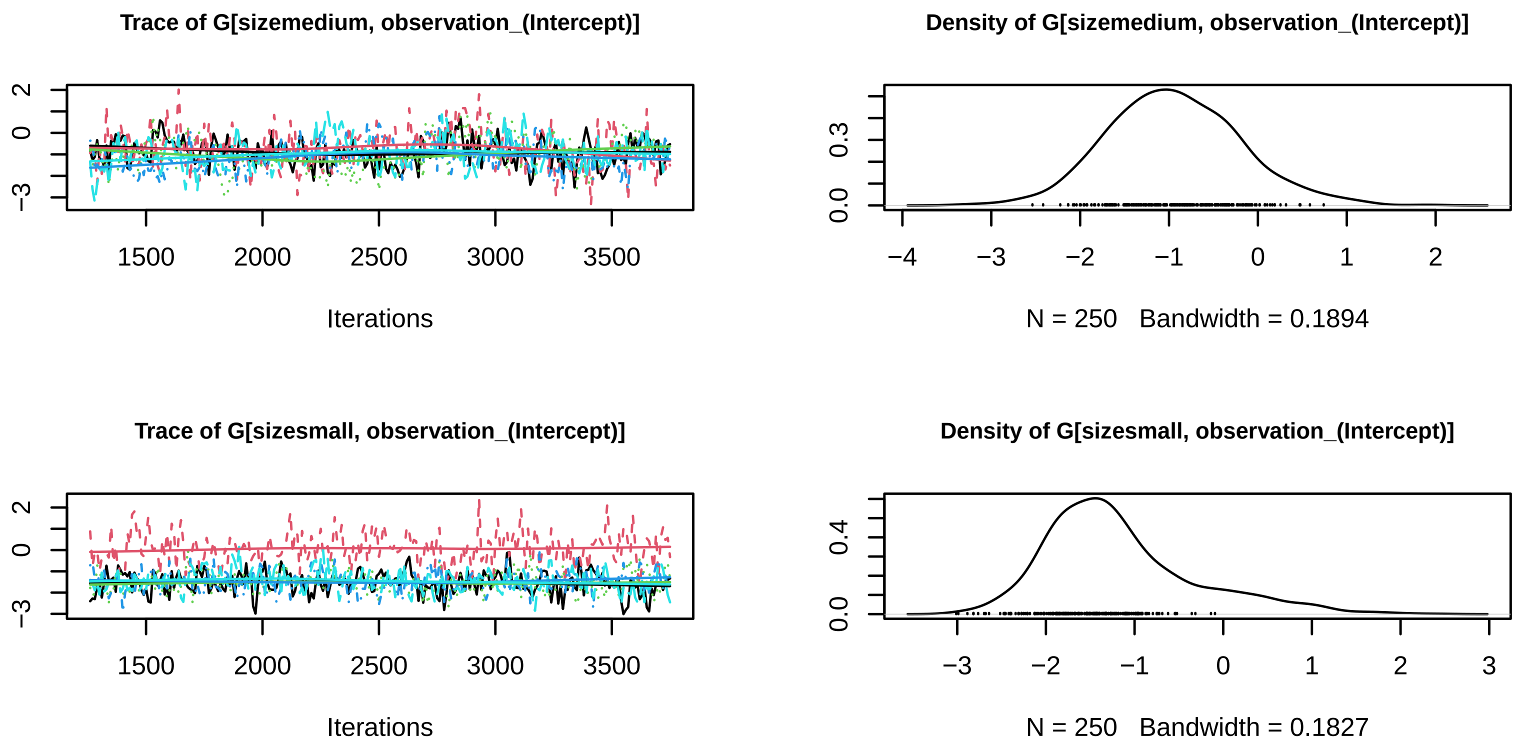
Trace of G[sizemedium, observation_(Intercept)]
Density of G[sizemedium, observation_(Intercept)]
Iterations
N = 250 Bandwidth = 0.1894
Trace of G[sizesmall, observation_(Intercept)]
Density of G[sizesmall, observation_(Intercept)]
Iterations
N = 250 Bandwidth = 0.1827

Figure S3: Mixing of the MCMC sampler for each of the Jsmm movement and observation parameters.

Table S1: Group identification, taxon, size, number of marked individuals, and total number of recaptures occured in Malaise traps

| ID | Taxon | Size | N marked individuals | Total n of recaptures |
|---|---|---|---|---|
| 1 | Aculeata | small | 12 | 0 |
| 2 | Auchenorrhyncha | small | 43 | 2 |
| 3 | Brachycera | small | 695 | 24 |
| 4 | Coleoptera | small | 113 | 6 |
| 5 | Coleoptera | medium | 2 | 0 |
| 6 | Heteroptera | small | 17 | 0 |
| 7 | Lepidoptera | large | 24 | 5 |
| 8 | Lepidoptera | medium | 90 | 1 |
| 9 | Lepidoptera | small | 871 | 43 |
| 10 | Nematocera | small | 140 | 8 |
| 11 | Nematocera | large | 5 | 5 |
| 12 | Nematocera | medium | 1 | 1 |
| 13 | Neuroptera | small | 16 | 1 |
| 14 | Parasitica | medium | 14 | 1 |
| 15 | Parasitica | small | 577 | 20 |
| 16 | Plecoptera | small | 1 | 0 |
| 17 | Sternorrhyncha | small | 4 | 0 |
| 18 | Symphyta | small | 16 | 0 |
| 19 | Symphyta | medium | 8 | 0 |
| 20 | Symphyta | large | 3 | 0 |

Table S2: Parameters estimates and convergence diagnostics of the MCMC approach used to sample the posterior distribution of the Malaise empirical data. The estimated parameters are $\beta_s^a$, $\beta_s^m$, $\beta_s^o$, where $s = 1, \cdots, 20$ represents the groups (Supplementary material Table 1.). The table shows the posterior mean, 2.5% and 97.5% posterior quantiles, the potential scale reduction factor (R-hat), the upper 95% confidence limit (Upper C.I.), and the Effective sample size (n.eff). Parameter estimates for the JSMM model are expressed in log form.

| Parameter | Mean | 0.025 Quantile | 0.975 Quantile | R-hat | Upper C.I. | n.eff |
|---|---|---|---|---|---|---|
| $\beta_1^a$ | 5.646 | 4.087 | 7.517 | 1.106 | 1.265 | 214.880 |
| $\beta_2^a$ | 5.198 | 3.102 | 8.421 | 1.490 | 2.213 | 168.908 |
| $\beta_3^a$ | 5.141 | 4.168 | 6.257 | 1.007 | 1.022 | 590.772 |
| $\beta_4^a$ | 5.050 | 3.768 | 6.592 | 1.033 | 1.091 | 333.465 |
| $\beta_5^a$ | 4.826 | 2.373 | 7.284 | 1.027 | 1.076 | 163.806 |
| $\beta_6^a$ | 5.809 | 3.498 | 8.259 | 1.030 | 1.083 | 239.016 |
| $\beta_7^a$ | 5.987 | 4.865 | 7.320 | 1.035 | 1.096 | 494.209 |
| $\beta_8^a$ | 6.343 | 4.514 | 8.257 | 1.089 | 1.230 | 231.795 |
| $\beta_9^a$ | 6.201 | 5.353 | 7.208 | 1.026 | 1.063 | 449.248 |
| $\beta_{10}^a$ | 5.347 | 3.187 | 7.310 | 2.677 | 5.089 | 363.227 |
| $\beta_{11}^a$ | 3.827 | 2.118 | 5.782 | 1.051 | 1.139 | 302.024 |
| $\beta_{12}^a$ | 4.256 | 1.914 | 6.766 | 1.156 | 1.390 | 156.798 |
| $\beta_{13}^a$ | 4.984 | 2.470 | 7.480 | 1.053 | 1.143 | 244.852 |
| $\beta_{14}^a$ | 5.133 | 3.072 | 7.293 | 1.042 | 1.110 | 225.116 |
| $\beta_{15}^a$ | 5.633 | 4.796 | 6.754 | 1.001 | 1.004 | 465.471 |
| $\beta_{16}^a$ | 5.610 | 2.924 | 8.120 | 1.059 | 1.156 | 293.695 |
| $\beta_{17}^a$ | 5.237 | 3.100 | 8.226 | 1.567 | 2.353 | 190.704 |
| $\beta_{18}^a$ | 5.986 | 3.538 | 8.407 | 1.069 | 1.179 | 154.278 |
| $\beta_{19}^a$ | 5.516 | 2.903 | 8.172 | 1.039 | 1.105 | 148.487 |
| $\beta_{20}^a$ | 5.328 | 2.979 | 7.663 | 1.014 | 1.043 | 153.726 |
| $\beta_1^m$ | -1.290 | -2.649 | 0.282 | 1.087 | 1.212 | 385.707 |
| $\beta_2^m$ | -2.451 | -4.279 | -0.617 | 1.698 | 2.614 | 270.222 |
| $\beta_3^m$ | -0.922 | -1.961 | -0.199 | 1.001 | 1.004 | 681.129 |
| $\beta_4^m$ | -0.959 | -2.513 | 0.167 | 1.007 | 1.021 | 291.390 |
| $\beta_5^m$ | -0.864 | -3.651 | 1.835 | 1.085 | 1.214 | 112.983 |
| $\beta_6^m$ | -1.054 | -3.449 | 1.259 | 1.028 | 1.075 | 266.679 |
| $\beta_7^m$ | -1.540 | -3.085 | -0.235 | 1.036 | 1.096 | 378.359 |
| $\beta_8^m$ | -0.661 | -3.439 | 1.759 | 1.085 | 1.209 | 98.999 |
| $\beta_9^m$ | -1.000 | -1.736 | -0.415 | 1.011 | 1.026 | 739.807 |
| $\beta_{10}^m$ | -2.072 | -3.481 | -0.670 | 1.647 | 2.524 | 291.750 |
| $\beta_{11}^m$ | -2.437 | -4.178 | -0.760 | 1.012 | 1.032 | 262.770 |
| $\beta_{12}^m$ | -1.766 | -4.401 | 0.273 | 1.078 | 1.194 | 129.143 |
| $\beta_{13}^m$ | -1.599 | -3.901 | 0.392 | 1.010 | 1.025 | 245.215 |
| $\beta_{14}^m$ | -1.399 | -4.218 | 1.037 | 1.069 | 1.178 | 89.608 |
| $\beta_{15}^m$ | -1.515 | -2.786 | -0.598 | 1.003 | 1.007 | 355.345 |
| $\beta_{16}^m$ | -1.400 | -3.773 | 0.999 | 1.010 | 1.018 | 269.848 |
| $\beta_{17}^m$ | -1.933 | -4.123 | 0.616 | 1.671 | 2.558 | 213.421 |
| $\beta_{18}^m$ | -1.061 | -3.456 | 1.300 | 1.075 | 1.194 | 130.675 |
| $\beta_{19}^m$ | -1.025 | -4.296 | 1.914 | 1.140 | 1.345 | 86.667 |
| $\beta_{20}^m$ | -1.692 | -4.194 | 0.748 | 1.096 | 1.241 | 125.004 |
| $\beta_1^o$ | -3.021 | -5.126 | -1.667 | 1.206 | 1.544 | 523.118 |
| $\beta_2^o$ | -1.873 | -4.864 | 3.776 | 5.386 | 9.526 | 282.432 |
| $\beta_3^o$ | -2.905 | -3.572 | -2.196 | 1.001 | 1.004 | 861.784 |
| $\beta_4^o$ | -2.515 | -3.642 | -1.414 | 1.036 | 1.099 | 451.498 |
| $\beta_5^o$ | -2.425 | -5.929 | -0.133 | 1.178 | 1.439 | 223.971 |
| $\beta_6^o$ | -3.511 | -5.824 | -1.543 | 1.030 | 1.063 | 448.830 |
| $\beta_7^o$ | -0.888 | -1.952 | 0.182 | 1.024 | 1.069 | 554.072 |
| $\beta_8^o$ | -2.513 | -4.713 | -0.952 | 1.301 | 1.708 | 203.574 |
| $\beta_9^o$ | -2.034 | -2.617 | -1.362 | 1.015 | 1.039 | 572.078 |
| $\beta_{10}^o$ | -0.755 | -2.917 | 4.725 | 9.041 | 16.538 | 421.146 |
| $\beta_{11}^o$ | 0.310 | -0.671 | 1.246 | 1.033 | 1.092 | 571.452 |
| $\beta_{12}^o$ | -0.762 | -2.631 | 1.270 | 1.137 | 1.327 | 206.049 |
| $\beta_{13}^o$ | -2.759 | -4.600 | -0.995 | 1.018 | 1.048 | 360.395 |
| $\beta_{14}^o$ | -2.513 | -4.537 | -0.847 | 1.040 | 1.080 | 267.015 |
| $\beta_{15}^o$ | -3.003 | -3.698 | -2.239 | 0.998 | 1.000 | 528.946 |
| $\beta_{16}^o$ | -2.889 | -5.597 | -0.332 | 1.048 | 1.054 | 310.620 |
| $\beta_{17}^o$ | -1.679 | -5.159 | 3.781 | 4.172 | 7.352 | 245.701 |
| $\beta_{18}^o$ | -3.753 | -6.187 | -1.708 | 1.033 | 1.053 | 275.990 |
| $\beta_{19}^o$ | -3.410 | -6.619 | -1.082 | 1.141 | 1.331 | 232.819 |
| $\beta_{20}^o$ | -2.454 | -5.286 | -0.352 | 1.211 | 1.518 | 262.120 |

Table S3: Descriptive statistics number of individuals per hectare that leads to expected number of individuals captured being one for each of the 20 groups.

| ID | Median | Mean | 1st Qu. | 3rd Qu. |
|---|---|---|---|---|
| 1 | 46241.0 | 146561.0 | 30792.0 | 74561.0 |
| 2 | 49060.8 | 79487.4 | 17889.0 | 102118.8 |
| 3 | 46346.0 | 48504.0 | 36451.0 | 57739.0 |
| 4 | 31088.0 | 36050.0 | 21646.0 | 43855.0 |
| 5 | 24030.0 | 3521000.0 | 11160.0 | 56580.0 |
| 6 | 80303.0 | 180666.0 | 39276.0 | 164584.0 |
| 7 | 6183.0 | 7007.0 | 4297.0 | 8651.0 |
| 8 | 28556.0 | 57940.0 | 16204.0 | 52522.0 |
| 9 | 19507.0 | 20097.0 | 15572.0 | 23752.0 |
| 10 | 18052.6 | 18364.0 | 10237.6 | 26372.7 |
| 11 | 1861.0 | 2139.0 | 1358.0 | 2615.0 |
| 12 | 5647.5 | 8796.3 | 2905.1 | 10133.1 |
| 13 | 37935.0 | 61742.0 | 22188.0 | 70945.0 |
| 14 | 29504.0 | 56544.0 | 16346.0 | 55581.0 |
| 15 | 50853.0 | 54011.0 | 39399.0 | 65826.0 |
| 16 | 45526.0 | 128756.0 | 19157.0 | 97814.0 |
| 17 | 35072.6 | 76718.9 | 10824.7 | 86063.4 |
| 18 | 97454.0 | 267373.0 | 50024.0 | 208998.0 |
| 19 | 69041.0 | 288132.0 | 32175.0 | 160286.0 |
| 20 | 26164.0 | 161994.0 | 12397.0 | 56738.0 |
